# Seasonal assembly of the phyllosphere fungal microbiome of a perennial grass is robust to nutrient addition

**DOI:** 10.1101/2025.05.08.652919

**Authors:** Elizabeth T. Green, Ignazio Carbone, Charles E. Mitchell

**Affiliations:** Biology Department, University of North Carolina at Chapel Hill, Chapel Hill, NC, USA; School of Plant Sciences, University of Arizona, Tucson, AZ, USA; Center for Integrated Fungal Research, Department of Entomology and Plant Pathology, North Carolina State University, Raleigh, NC 27695, USA

**Keywords:** phenology, microbiome, phyllosphere, fungal community, community recruitment

## Abstract

The leaf microbiome plays an important role in plant health and defense. Despite its importance, how the assembly of the leaf microbial community is modified by environmental conditions such as nutrient availability remains relatively uninvestigated. Soil nutrient availability may shift the outcome of microbial interactions within a host individual or influence the pool of microbes across the plant community. We hypothesized that leaf microbial diversity would increase across the season as leaves collect additional taxa, and that this seasonal assembly would be sensitive to nutrient addition. To assess this, we tracked the assembly of the fungal phyllosphere microbiome of the grass tall fescue (*Lolium arundinaceum*) in old-field vegetation over the growing season and experimentally tested whether the seasonality of the microbiome was modified by experimental addition of soil nutrients. Fungal diversity (Shannon diversity index, richness, and evenness) increased early in the season, with most metrics saturating before the end of the season. Community composition as measured by Bray-Curtis dissimilarity also shifted over the early and mid-growing season. Phylogeny-based machine-learning identified fungal lineages that were abundant in different seasons, linking seasonal community shifts to their evolutionary context. Nutrient addition was less important than time of season, but still significantly altered community composition and interacted with time to influence richness, with lowest richness in the low nutrient addition plots early in the season. The clear seasonality of the microbiome provides support for a dynamic phyllosphere microbiome, suggesting further studies manipulating fungal recruitment over the season. Furthermore, it highlights the robustness of seasonal assembly to variation in nutrient availability.

## Introduction

The leaf phyllosphere, which includes the leaf surface and interior, supports a dense and diverse community of bacteria and fungi, which interact with their hosts and each other to alter disease and herbivory (Berg & Koskella, 2018; Busby et al., 2016; Humphrey et al., 2014; Ritpitakphong et al., 2016) as well as nutrient cycling (Fürnkranz et al., 2008). A leaf’s microbial community is shaped by many factors including the soil microbial community, surrounding plant species, vegetation density, and seasonal environmental and abiotic fluctuations (Borer et al., 2013; Grady et al., 2019; Liu et al., 2017; Seabloom et al., 2023; Trivedi et al., 2020). In the phyllosphere, the structure of fungal community is known to shift dynamically on multiple temporal scales including seasonally (Argiroff et al., 2024; Boutin & Laforest-Lapointe, 2025; S. Wang et al., 2025). For most species in temperate zones, the leaf microbiome is reestablished each year and follows a successional trend. At the beginning of the growing season, many plants begin putting up new shoots and leaves. The initial, early season microbial community on those leaves is strongly related to the soil, leaf litter, and rhizosphere microbial community (C. Gao et al., 2020; Kong et al., 2019). As the season progresses, the microbial community changes in response to the environmental conditions and the influx of new microbes (Debray et al., 2022). Later in the season, the leaf community shifts to contain more leaf-specific taxa and to mirror more closely that of surrounding plants (F.-L. Gao et al., 2019).

In addition to transient microbial taxa that are sporadically present throughout the season, recent research efforts have attempted to characterize the core microbial community (Custer et al., 2023; Howe et al., 2023; Shade & Handelsman, 2012; Shade & Stopnisek, 2019). The core community is a set of frequently occurring and abundant taxa (Cantoran et al., 2023; Hernandez-Agreda et al., 2017; Shade & Handelsman, 2012; Umaña et al., 2017) that can be key players in organismal processes, including by increasing abiotic stress tolerance, promoting growth, and protecting against foliar pathogens (Grady et al., 2019; Howe et al., 2023; Shade & Handelsman, 2012; Trivedi et al., 2020). Environmental nutrients are one of the abiotic factors that can affect the plant microbiome. Humans have increased the input of nutrients to terrestrial systems through fertilization and burning of fossil fuels (Keller et al., 2023; Steffen et al., 2015; Veresoglou et al., 2013), so decoding the nuanced effects of nutrients across organizational levels is imperative. Nutrient addition can alter the plant tissue nutrient concentrations and thus the resources available to microbes, which can alter the microbiome diversity and composition (Firn et al., 2019; Lumibao et al., 2019; Oono et al., 2020; Tellez et al., 2022). Additionally, host nutrient availability can alter the host immune response to different microbes and the outcomes of nutrient-mediated microbial interactions (Borer et al., 2013; Lacroix et al., 2014; Smith et al., 2009). Given these diverse potential effects of nutrient addition on the leaf microbiome and the seasonally dynamic assembly of leaf microbiota (Grady et al., 2019; Howe et al., 2023), we need to understand the influence of nutrients on seasonal assembly of the leaf microbiome.

To assess the impact of soil nutrients on the seasonal assembly of the leaf microbiome, this study focused on the fungal community in the phyllosphere of tall fescue (*Lolium arundinaceum*). While the seasonal dynamics of the leaf microbiome have been previously characterized across crop and wild species, there are still large gaps in our knowledge of those seasonal dynamics, including how the seasonal assembly of transient microbes vs. the core community is modulated by soil nutrient abundance (Dale & Newman, 2022; Debray et al., 2022; C. Gao et al., 2020; Grady et al., 2019; Trivedi et al., 2020). We built on a previous study of the core fungal microbiome of tall fescue (Dale & Newman, 2022) to experimentally test how the seasonal assembly of the core and complete fungal microbiomes is modulated by soil nutrients. Additionally, we integrate functional analysis of fungal trophic mode with seasonal dynamics and apply machine learning to phylogenetic trees to identify specific taxa and trophic modes associated with early- and late-season communities. Together, this study addresses the effect of soil nutrient addition on the function of rare and core taxa in the leaf microbiome across a growing season to address three hypotheses at the intersection of microbial seasonal assembly and nutrient abundance.

First, we hypothesized that we would see an increase in fungal community diversity throughout the season. Second, we hypothesized that early in the season a core microbial community would establish that was consistent across replicate plots, then over the season, replicate plots would diverge in taxonomic community composition as different plots accumulated different microbial taxa, increasing total richness and expanding the complete community. Third, we hypothesized that soil nutrient addition would alter the functional composition of the phyllosphere community by shifting within-host microbial interactions and favoring fungi that are better able to exploit increasing leaf nutrient availability, including pathotrophic taxa. To test these three hypotheses, this study evaluated the seasonal leaf microbiome assembly in a field setting and experimentally tested the influence of soil nutrients on the leaf microbiome.

## Methods

### Field setup

The experiment was conducted across a single growing season at Widener Farm (Duke Forest Teaching and Research Laboratory). Widener Farm is an 8-ha old field located in Orange County, North Carolina that was last cultivated in 1996 (Heckman et al., 2016). The site is dominated by perennial grass and shrub species, including tall fescue (*Lolium arundinaceum*). Tall fescue supports a diverse assembly of foliar, fungal pathogens that cause epidemics of several diseases over the growing season, including anthracnose, brown patch, and rust (Grunberg et al., 2023; Halliday et al., 2019).

In early April of 2021, we primed and germinated tall fescue seed collected from Widener Farm in 2018 by soaking seeds in water for six hours and then left them out to dry overnight. We sprinkled the seed onto moist vermiculite and left them in a closed Tupperware container for 10 days. After that, we haphazardly selected seedlings to transfer to individual pots filled with MetroMix 360 soil. In the greenhouse, we watered the seedlings three times per week for eight weeks at which point plants were hardy enough to be moved outside.

Meanwhile, we staked out 72 1m x 1m plots at Widener Farm in a 6 x12 array. On May 2, 2021, we sprayed every other plot with Roundup ® 360 and the following week covered the sprayed plots with landscaping fabric to kill all above ground vegetation. This treatment produced a checkerboard pattern of 36 sprayed and 36 unsprayed plots (Figure S1). Over three days from May 31 to June 2, 2021, we removed the landscaping fabric and transferred the seedlings from the greenhouse to the sprayed plots. We planted 12 seedlings in each plot in a 3 x 4 grid.

In a randomized blocked design with three spatial blocks of 12 plots each, we randomly assigned each plot one of three treatments of 10-10-10 N-P-K Meherrin Fertilizer (no nutrients, low nutrient addition (5 g/m^2^), high nutrient addition (10 g/m^2^)). This yielded four plots of each nutrient treatment in each block. In October 2021, we mowed the plots, then treated them with their assigned nutrient treatments before letting them sit undisturbed over the winter. In September of 2021-2023, we surveyed the plant community by estimating percent area cover of all plant species within a 0.5m × 0.5m square in the center of each plot. We detected no effect of nutrient addition on either Shannon Diversity or Bray-Curtis dissimilarity of the plant community (Table S1-S2), so we did not consider the broader plant community in our analyses of the foliar fungal community on tall fescue.

### Sample Collection and Processing

Starting on April 29, 2022, we collected samples from the oldest leaf of three tall fescue tillers per plot, selected to be equally spaced throughout the plot, to assess the fungal microbiome of the fescue in the plots. If the oldest leaf was more than 50% chlorotic or necrotic, we collected from the second oldest leaf instead. Using an office hole punch sterilized with 70% ethanol between samples to prevent the transmission of disease between plants (Wagner et al., 2020), we collected three hole-punches of leaf tissue evenly spaced across the leaf. Hole-punches were taken of healthy tissue or tissue that was partially necrotic or diseased, but not of fully chlorotic or necrotic tissue. Leaf tissue samples were kept on ice and transported back to the University of North Carolina at Chapel Hill, about 20 minutes away, where, on the same day, we surface rinsed the samples by adding 1mL of sterile water to each sample and vortexing for 10 seconds to remove any soil or loosely associated microbes before removing the water and freezing the samples at -80C (Wagner et al., 2020). To retain epiphytes, samples were not surface-sterilized. Samples were collected on four additional sample days approximately 6 weeks apart (June 7, July 15, August 23, and October 4, 2022) for a total of 180 samples.

We ground the frozen leaf tissue samples in a ball mill (Retsch Mixer Mill) and extracted DNA with the DNeasy Power Soil Pro Kit (Qiagen) following the manufacturer protocol. We used between 0.2-0.4g of frozen, ground leaf material per DNA extraction. We completed DNA amplification and library preparation and barcoding using the Quick-16S NGS Library Prep Kit (Zymo Research) and substituted locus-specific ITS primers (ITS1F: 5-CTTGGTCATTTAGAGGAAGTAA -3; ITS2: 5-GCTGCGTTCTTCATCGATGC -3) for the provided 16S primers. Library preparation was completed on two 96-well plates with a negative control of sterile water on each plate (Geyer et al., 2024). . The negative controls did not amplify with qPCR. Five samples were randomly selected as duplicate controls between plates, yielding 185 samples to sequence after the removal of the negative controls. Samples were submitted to the High Throughput Sequencing Facility (University of North Carolina at Chapel Hill) where they tested the sample quality (plate 1: 51ng/μl, 392 base pairs; plate 2: 40.4 ng/μl, 373 base pairs) and sequenced the samples with a NextSeq 2000 P1 (Illumina) paired-end run.

### Data Analysis

ITS sequence files were imported, demultiplexed, and denoised with the DADA2 plugin (Callahan et al., 2016) to Qiime2 (Bolyen et al., 2019). We trimmed the primers from the reads but did not otherwise truncate reads to a fixed length. Reads were merged into an amplicon sequence variant (ASV) table, and we removed chimeric ASVs, also with Qiime2. The representative sequences were then phylogenetically placed on a six-locus fungal reference tree Fungi_v3 (T-BAS accession 4D2ZPWQH) (Carbone & White, 2025) with the Tree-Based Alignment Selector Toolkit (T-BAS v2.3) (Carbone et al., 2019). The power of phylogenetic methods largely rests on the reliability of the phylogeny, which can be uninformative if based on a single locus. T-BAS reduces uncertainty in the phylogenetic tree because microbiome sequences are placed on well resolved multilocus reference trees. Final taxonomic assignments were based on comparing placement results from T-BAS with assignments from UNITE Version 10.05.2021 (Põlme et al., 2020), RDP classifier Version 2.13 (Q. Wang & Cole, 2024), and BLAST of ASVs against the NCBI non-redundant database (Camacho et al., 2009)(Table S3). We assigned trophic mode to the placed ASVs using FunGuild at the genus level (Nguyen et al., 2016).

We removed ASVs that were not assigned to the kingdom fungi and further removed “nonreproducible” ASVs that were not observed at least 25 times in each of at least five samples (Lundberg et al., 2012; Wagner et al., 2020). We finally removed samples with <2000 usable reads to remove outlier samples from further analyses. This filtering reduced the data set to 359 ASVs and excluded 21 of the 185 samples. We compared the read depth of the duplicate samples to confirm limited variation between the 96-well plate preparations and removed samples of each duplication at random. The number of reads remaining per sample was saved to use as a representation of sampling effort in further analyses. All further analyses treated ASVs as taxa. We variance normalized the count data (DESeq package) (Love et al., 2014) for ordination analyses.

To identify the core fungal community, we followed Shade and Stopnisek (Shade & Stopnisek, 2019), an approach based on each taxon’s contribution to community composition, measured by Bray-Curtis dissimilarity. First, to rank taxa as candidates for inclusion in the core, we calculated an index for each taxon that integrated its temporal consistency and spatial occupancy. We calculated the spatial occupancy of each taxon as the proportion of plots that the taxon was present in, for each of the five survey dates, yielding five values of occupancy. We calculated the temporal consistency of each taxon as 1 if present in all plots on a survey date or 0 if not, for each of the five survey dates, yielding five values of consistency. The index value for each taxon was calculated as:

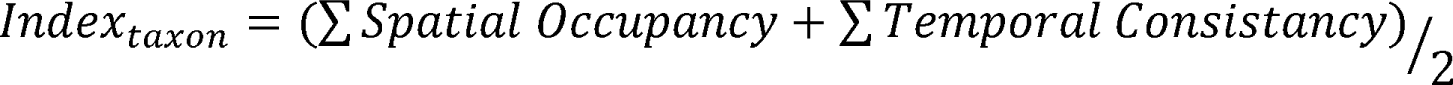

Considering both occupancy and abundance allowed us to identify taxa that persisted over time through environmental disturbance events, such as rain and intense heat, as well as spatially across a population of plants. To define core membership based on contribution to variation in community composition, we calculated the mean Bray-Curtis dissimilarity for the complete data. Next, in an iterative process starting with the taxa in the first index group (i.e. the greatest index value), we calculated the mean Bray-Curtis (BC) dissimilarity for the subset and determined the proportion of community dissimilarity explained by the subset of the complete community (1-(BC_core_/BC_all_)). We then proceeded in descending order to the taxa in the next lower index subset, each time adding those additional taxa to the taxa in the previous subset (Figure S2). We selected the cutoff for core taxa as where adding the next occupancy group explained a less than 0.03 difference in proportion of community dissimilarity from the complete community dissimilarity (Shade & Stopnisek, 2019). We determined the 0.03 cutoff to best describe the core community, but not the rare or transient taxa, by comparing the R^2^ values of the Bray-Curtis dissimilarly analysis with the cut off at 0.01 (R^2^ = 0.498), 0.02 (R^2^ = 0.511), 0.03 (R^2^ = 0.528), and 0.04 (R^2^ = 0.512).

As metrics of alpha diversity, we calculated Shannon diversity index and taxa richness of the complete and core communities of each sample. We then calculated evenness as the Shannon diversity index divided by the natural log of richness for both communities. We used mixed-effect models with fixed effects of sampling depth (number of reads per sample), date (as a continuous variable), nutrient addition, and the date × nutrient-addition interaction (lme4 package) (Bates et al., 2015). We nested plot within spatial block as random intercepts to account for any spatial correlations (Wagner et al., 2020). We also log transformed both metrics to reduce heteroscedasticity. We modeled each metric as a linear and quadratic function of sampling date and compared delta AIC (linear - quadratic, positive value indicates that the quadratic model is a better fit and negative value indicates that the linear model is a better fit) values to determine which best fit the data. We also analyzed the richness of each trophic mode of the complete and core communities as they interacted with survey date and nutrients. We performed a multivariate ANOVA with richness of each trophic mode as the dependent variables and fixed effects of sampling depth and the interaction between date and nutrient addition. .

To assess species composition, we conducted a Bray-Curtis dissimilarity analysis between treatments and sample dates using the variance-stabilized count data for both the complete and core communities. We ordinated the Bray-Curtis distance matrix to visualize the community relationships (vegan package) (Oksanen et al., 2022). We then ran a factorial PERMANOVA (vegan, adonis2) to model the Bray-Curtis dissimilarity values against the interaction of nutrient treatment and survey date and included sampling depth as a fixed effect. The model was grouped by plot nested within block to account for spatial correlations in the sampling. We calculated the dispersion of each sample as the distance from the centroid for that survey date and nutrient treatment combination (vegan, betadispr). We then modeled the distances as a linear mixed effect model with a factorial of date and nutrient addition, with sampling depth as an additional fixed effect and plot nested within block as random intercepts.

Finally, we employed two methods to investigate whether there were specific taxa correlated with nutrient addition or survey date. First, we conducted an indicator species analysis (indicspecies package) (Cáceres & Legendre, 2009) to analyze the taxa richness by the interaction of nutrient addition and survey date. Second, we applied machine learning to explore which taxa could discriminate between (A) the high vs. low nutrient treatments, and (B) the first vs. last seasonal sampling dates, using PopPhy-CNN, within T-BAS. T-BAS tracks the common ancestry of microbes because common ancestry can be both informative (e.g. in exploring correlations among traits or taxa abundances) and a source of confounding variation (e.g. in regression analysis of single and multiple traits and variables) if not dealt with in microbiome studies (Washburne et al., 2018). Moreover, phylogenetic-based methods can be essential to predictive modeling of microbiome data (Xiao et al., 2018). PopPhy-CNN applies convolutional neural networks to phylogenetic trees to identify microbial taxa that can predict system features such as an experimental treatment, location, or time (Reiman et al., 2020). To implement this approach, we first placed all ASVs on a reference tree comprising over 2,300 fungal taxa, inferred from multilocus sequence data (Fungi_v3; T-BAS Accession 4D2ZPWQH) (Carbone & White, 2025). This reference tree was expanded from a previously published six-gene phylogeny of 199 fungal taxa (James et al., 2006). As well as assigning many ASVs to taxonomic groups, this phylogenetic framework allowed us to retain both assigned and unassigned taxa for the CNN model. Phylogenetic placements were then combined with ASV count data to train the CNN models. Each model was trained using 10-fold cross validation, an L2 regularization rate of 0.001, and a kernel size of 4 (width) × 3 (height).

In our experiment, we developed two models, each of which predicted (discriminated between) two binary classes: 1) high nutrient addition with 57 samples vs. no nutrient addition (unfertilized) with 60 samples, and 2) early season (survey on 4/30/2022) with 36 samples vs. late season (survey on 10/4/2022) with 36 samples. Full taxonomic trees were pruned based on the ASVs in each dataset. For each binary class (e.g. early vs. late season), we identified taxa and lineages that discriminated between the two feature states based on feature scores derived from the first convolutional layer, which offers the highest resolution (Reiman et al., 2020). Feature scores were calculated for ASVs and for genera, then all feature scores were ranked together. Internal tree nodes with larger feature scores than their children indicate that the feature was more discriminative at a higher taxonomic level. Feature scores exceeding the 90th percentile (i.e., scores with absolute value > 1.0) were considered the most informative and examined further. Phylogenetic trees with annotated nodes and edges based on calculated feature scores were visualized using Cytoscape Version 3.10.3 (Shannon et al., 2003).

## Results

The final database after filtering included 359 taxa (ASVs) across 164 samples. The total number of sequence reads per sample (sampling depth) ranged from 5,802 to 448,078 with a median of 86,241 (Figure S3).

Based on the approach of Shade and Stopnisek (Shade & Stopnisek, 2019), the core community comprised 54 taxa (Table S4) that were found in the first 28 ranked index values of 149 total values. These core taxa accounted for 86.6% of the total reads.

### Hypothesis 1: Phyllosphere fungal microbiome increased in diversity over the growing season but was robust to nutrients

Metrics of alpha diversity were highly structured by date for both the complete and the core community. We first tested whether the relationship of each diversity metric to time of season was better fit by linear or quadratic equations, based on AIC. Richness of the complete community increased linearly across the season (delta AIC = -38.40). Shannon diversity index, evenness, and richness of the core community were modeled best by a quadratic equation (delta AIC; complete Shannon = 18.09; core Shannon = 30.96; complete evenness = 9.55; core evenness = 20.24; core richness = 6.22). Thus, all metrics of diversity, except for richness of the complete community, increased fastest early in the season before leveling out later in the season.

For both the complete and core communities, the Shannon diversity index (complete: conditional R^2^ = 0.81, F_1,164_ = 127.25, p < 0.001, Table S5; core: conditional R^2^ = 0.73, F_1,164_ = 131.21, p < 0.001, Table S6; Figure 1), richness (complete: conditional R^2^ = 0.76, F_1,164_ = 379.92, p < 0.001, Table S7; core: conditional R^2^ = 0.68, F_1,164_ = 107.57, p < 0.001 Table S8; Figure 2), and evenness (complete: conditional R^2^ = 0.73, F_1,164_ = 105.83, p < 0.001, Table S9; core: conditional R^2^ = 0.70, F_1,164_ = 116.95, p < 0.001, Table S10; Figure 3) were significantly predicted by the survey date. Of the three metrics of alpha diversity analyzed, only foliar fungal richness was sensitive to the interaction of survey date and nutrient addition for both complete and core communities with the low nutrient addition plots having the lowest taxa richness and lowest rate of richness increase across the survey dates (date * nutrients; complete: F_2,164_ = 172.49, p < 0.001, Table S7; core: F_2,164_ = 4.74, p = 0.010, Table S5).

**Figure 1.**
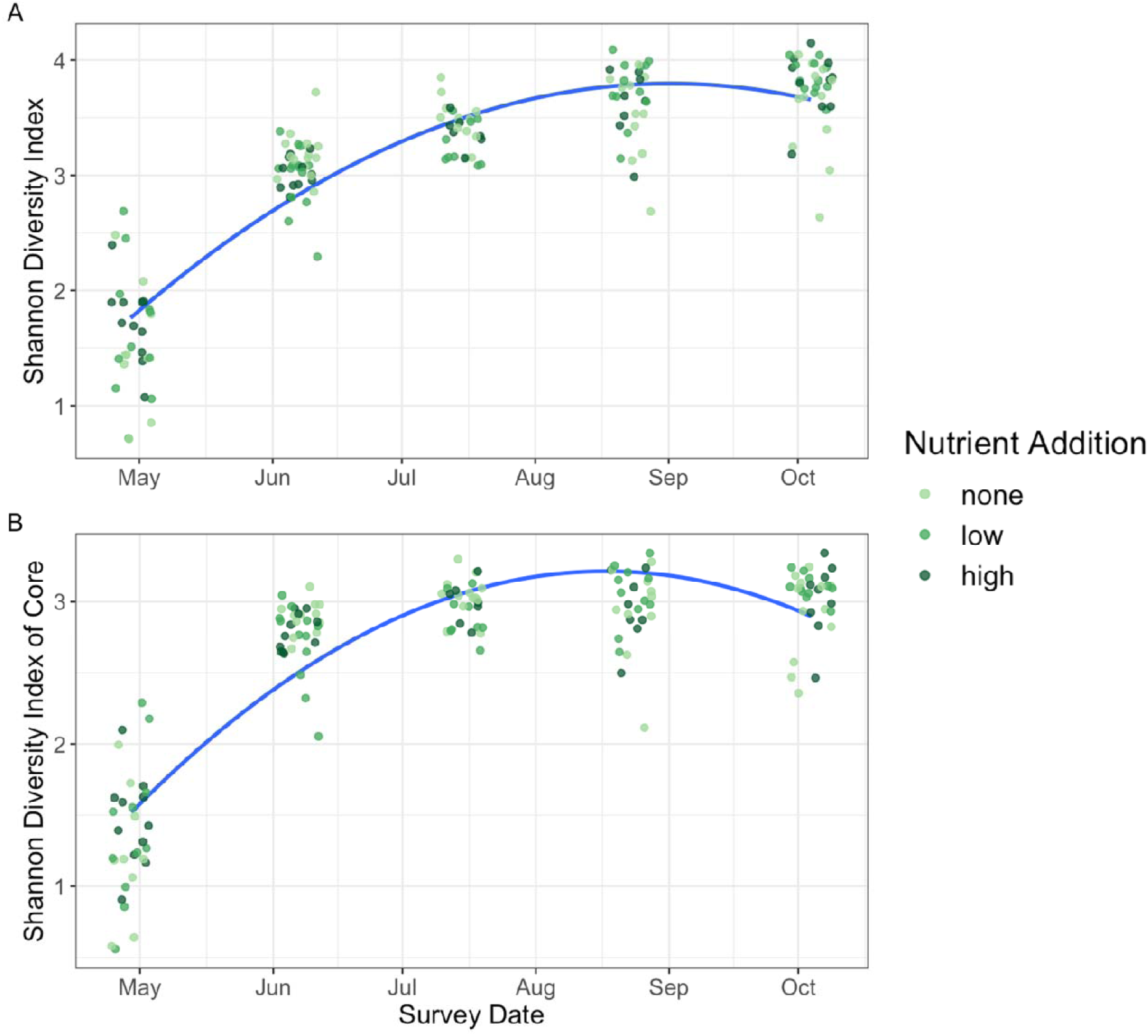
Shannon diversity index of complete (A) and core (B) communities across survey dates with quadratic fit lines. For both, diversity increased early in the season, then leveled off. Colors represent the nutrient addition treatment.

**Figure 2.**
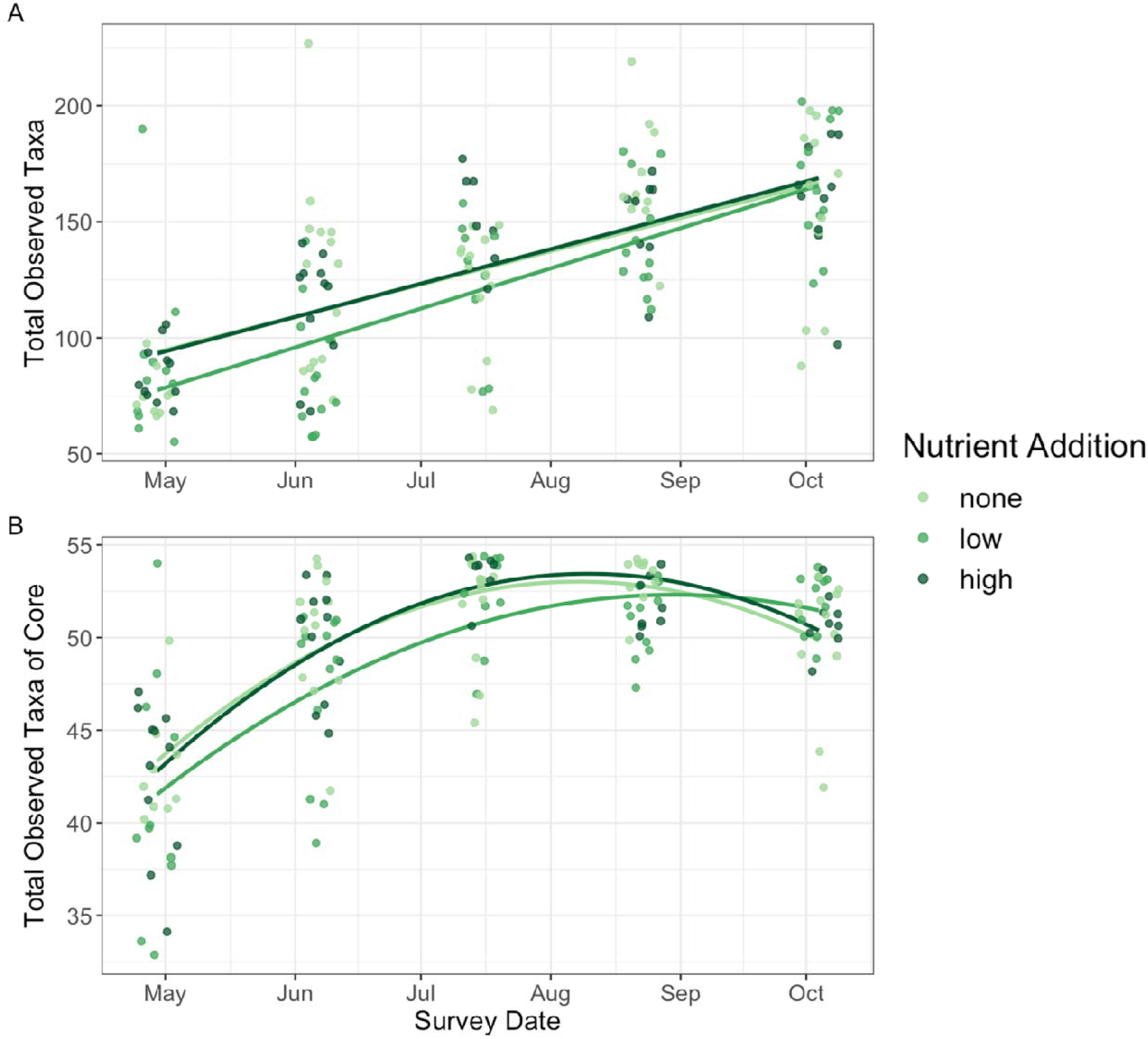
Taxa richness across survey dates for the complete (A, linear) and core (B, quadratic) communities. For both, there was a statistically significant interaction between nutrient addition and survey date, with the lowest taxa richness in the low nutrient addition plots early in the season. Color represents the nutrient addition treatment.

**Figure 3.**
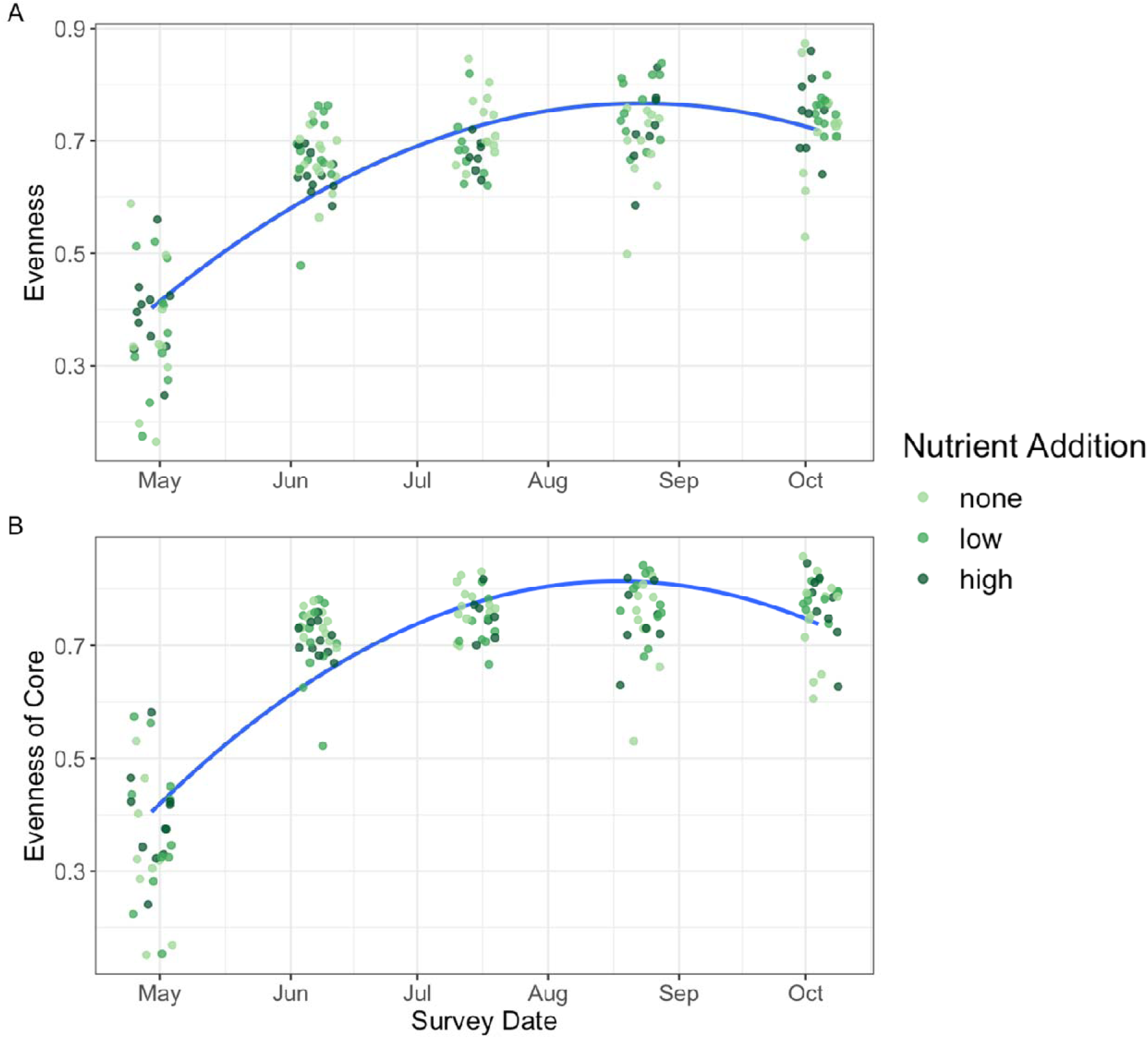
Taxa evenness across survey dates for the complete (A) and core (B) community with quadratic fit lines. For both, evenness increased early in the season, then leveled off. Color represents the nutrient addition treatment.

### Hypothesis 2: A core microbial community established early in the season and interacted with nutrient addition to diverge as plots increased in richness across the season

When assessing the composition of the complete community, there was a clear trend across the sampling dates as the Bray-Curtis dissimilarity shifted from early to late-season and the ordination of communities moved across two axes (F_1,164_ = 48.96, p < 0.001, Table S11, Figure 4A). Survey date explained 21.3% of the variation between communities. Community dispersion (or the distance to the centroid) of the complete community also increased across the season (date: F_1,164_ = 68.69, p < 0.001; Table S12, Figure 5A). Survey date explained a greater amount of the variation of the core community, 35.2% (F_1,164_ = 113.21, p < 0.001, Table S13, Figure 4B), than when looking at the complete community. However, dispersion of the core community was level across the survey dates (date: F_1,164_ = 10.32, p = 0.002; Table S14, Figure 5B).

**Figure 4.**
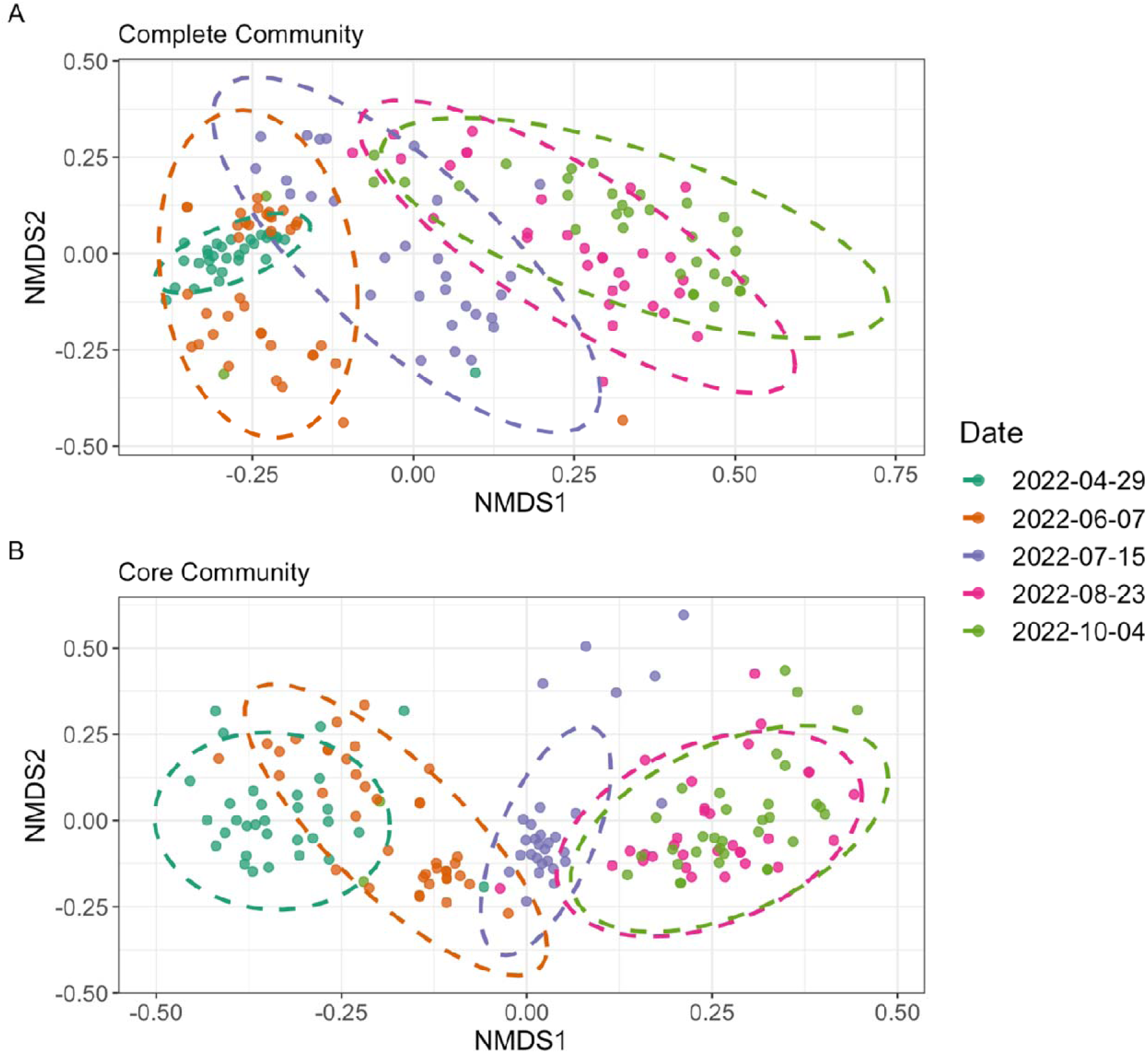
NMDS based on Bray-Curtis dissimilarity of the complete (A) and core (B) fungal taxa in each plot, grouped by survey date. 95% confidence ellipses are shown for each date.

**Figure 5.**
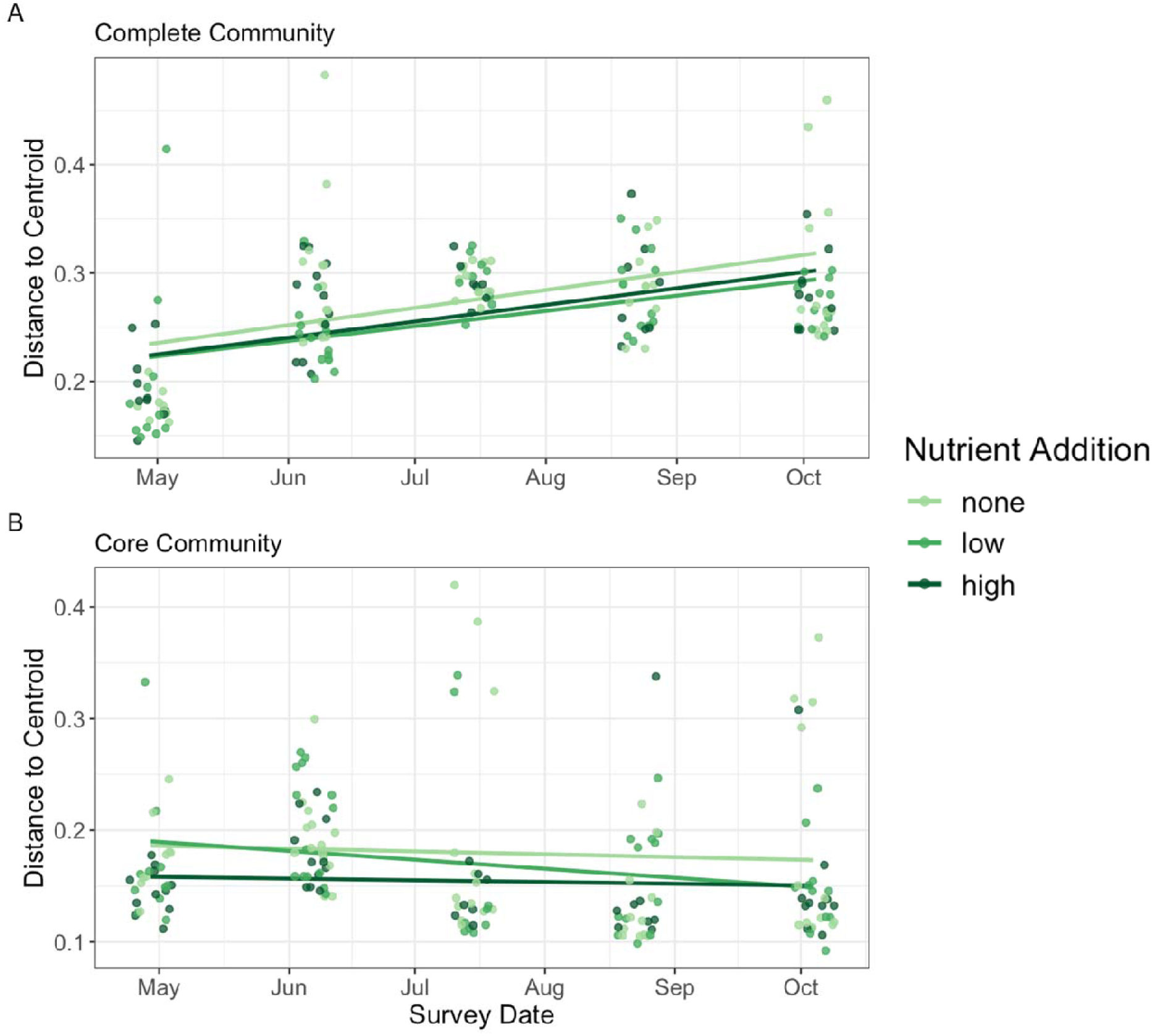
Community dispersion as measured by the distance to centroid of each survey date for the NMDS based on the Bray-Curtis dissimilarity of the complete (A) and core (B) fungal taxa in each plot. Color represents the nutrient treatment.

Nutrient addition had a significant effect on Bray-Curtis dissimilarity, but explained only 1.9% of the variation (F_2,164_= 2.28, p < 0.001, Table S11, Figure S4A). The interaction between date and nutrient addition trended towards significant but was not significant (date * nutrients: F_2,164_ = 1.52, p = 0.083). Community dispersion also interacted with nutrient addition as the unfertilized plots increased in dispersion more quickly than the low or high fertilizer plots (conditional R^2^ = 0.37; date * nutrients: F_2,164_ = 32.38, p < 0.001 Table S12, Figure 5A). Nutrient addition explained a smaller, 0.8%, but still significant, amount of the variation of the core community (F_2,164_ = 1.43, p = 0.020, Table S13, Figure S4B), and did not interact with survey date (nutrients * survey date; F_2,164=_ 1.40, p = 0.268). Dispersion of the core community interacted with nutrient addition with low nutrient plots decreasing in dispersion across the season while dispersion in unfertilized and high nutrient plot remained stable (conditional R^2^ = 0.47; date * nutrients: F_1,164_ = 5.33, p = 0.006, Table S14, Figure 5B).

### Hypothesis 3: Fungal trophic modes and predictive taxa differed between early- and late-season communities, but not nutrient addition

Distribution of fungal trophic modes shifted across the season with the later-season communities becoming dominated by taxa that were categorized as saprotrophs (date; complete: F_1,162_ = 9.61, p < 0.001, Table 15; core: F_1,162_ = 7.003, p = <0.001, Table S16; Figure 6). We did not find any effect of nutrient addition on the richness of trophic modes across the season (nutrients; complete: F_2,162_ = 0.65, p = 0.802, Table S15; core: F_1,162_ = 0.968, p = 0.461, Table S16; Figure 6).

**Figure 6.**
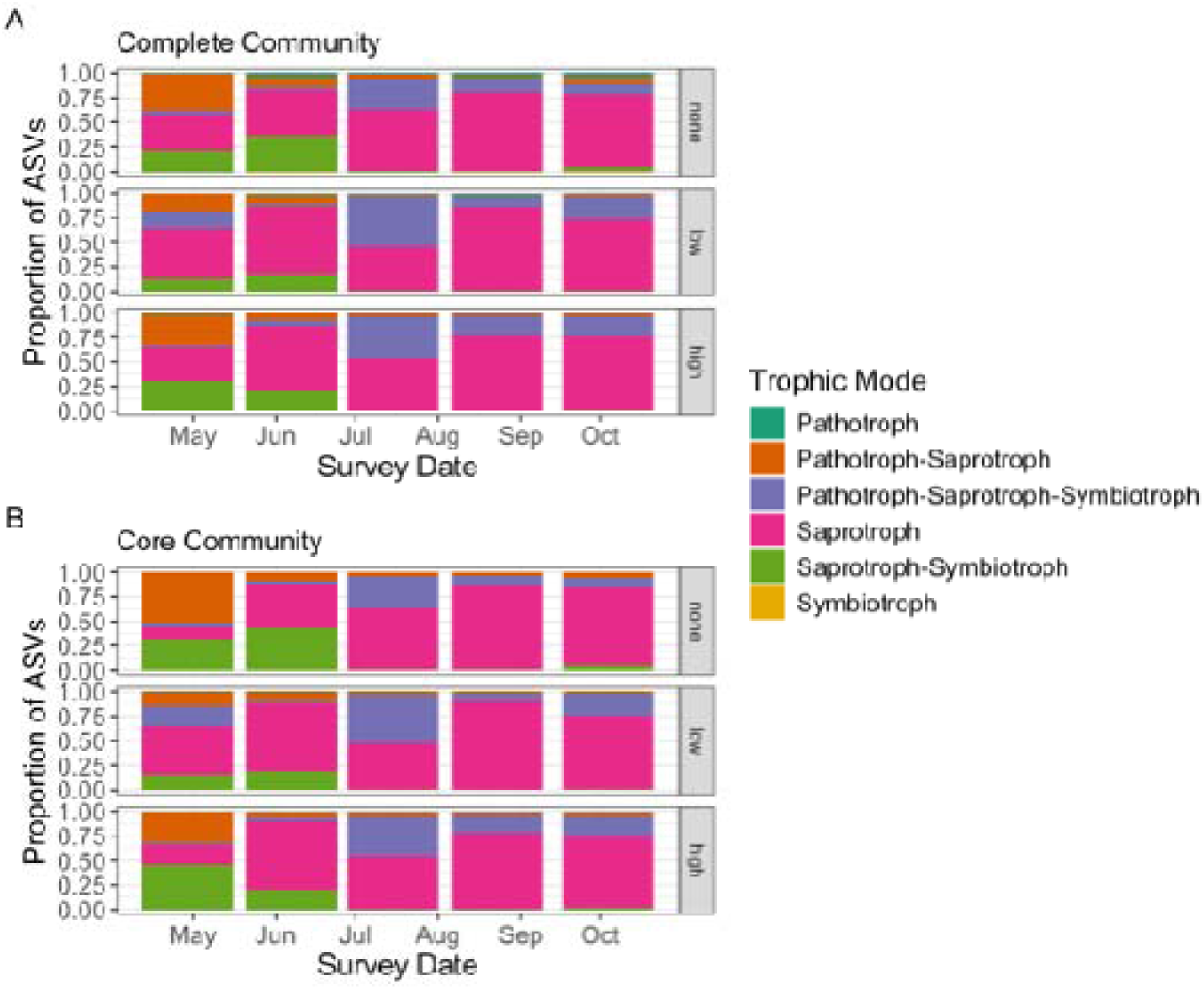
Proportion of ASVs assigned to each trophic mode for the complete (A) and core (B) communities. Proportions are calculated by survey date and nutrient treatment (none, low, and high).

The indicator species analysis identified no taxa as significantly associated with any of the survey dates or nutrient addition treatments. The convolutional neural networks were also not able to identify fungal lineages that predicted nutrient addition treatment (unfertilized feature score = 0.95, high nutrient addition feature score = -0.49), but the neural networks did identify fungal lineages that predicted survey date (early season feature score = 1.69, late season feature score = -1.70). There were 15 highly predictive taxa for early season communities and 22 highly predictive taxa of late season communities (Table S17; Figure S5). *Cladosporium* accounted for the top three predictive lineages and the top predictive taxon (ASV 40) for early season communities, while *Leucosporidium* had the greatest number of lineages (5) predictive for early season communities (Table S17). In late season communities, *Bulleribasidium* had the greatest number of predictive lineages (4) and the most predictive taxon (ASV 57).

## Discussion

There was a clear increase in community diversity across the seasonal surveys, in support of our first hypothesis. All three metrics of alpha diversity (Shannon diversity index, species richness, and species evenness) displayed increasing diversity early in the season for both the complete and the core microbiomes. Shannon diversity and evenness increased quadratically for both sets of communities with the fastest increase occurring early in the season before leveling off. Species richness increased quadratically for the core community, but richness of the complete community increased linearly across the season, suggesting that the complete and core microbiomes added taxa at different rates throughout the season. We identified a core community of 54 taxa. The remaining 305 taxa were transient and together contributed only 3% to the dissimilarity of the complete community (Shade & Stopnisek, 2019). This core community suggests that there is a consistent collection of fungi in the phyllosphere of tall fescue that comprises of a dominant set of taxa throughout the season, lending support to our hypothesis that the core complete communities add taxa at different rates.

One way to interpret the difference in richness trajectory between the complete and core communities is through the lens of community succession of the phyllosphere (Copeland et al., 2015; Debray et al., 2022; Grady et al., 2019; Howe et al., 2023). Early in the spring, tall fescue plants begin to put up new tillers and collect microbes from the soil and surrounding environment. Some fungi may overwinter in the root tissue and be vertically transmitted to the new tillers or in the soil but given the low species richness early in the season, this seems to be the minority of taxa (Kong et al., 2019; Otten et al., 2001). The new tillers begin recruiting microbes and soon have collected their community of core taxa. Throughout the rest of the season, more transient fungi continue to colonize the leaves and the phyllosphere shifts from a lower diversity, more homogenous early-successional community to a diverse and varied late-successional community. Once the core taxa became established, it persisted for the remainder of the season resulting in quadratic growth (Copeland et al., 2015; Grady et al., 2019). However, the complete community recruited taxa at a consistent rate throughout the season resulting in linear growth. As taxonomic diversity shifted over the season, so did community composition, as measured by Bray-Curtis dissimilarity of both core and complete communities. Furthermore, the dispersion of samples as measured by their distance from the centroid NMDS value of each survey date increased across the season for the complete community and did not shift substantially over the season for the core community.

While the diversity of both the core and complete communities increased from the early-successional to late-successional communities, the diversity of trophic modes decreased, with saprotrophs dominating the late-successional communities. Early-successional communities had a greater proportion of both symbiotrophs and pathotrophs, suggesting that the fungi overwintering in plant tissue may include both beneficial and pathogenic taxa. The accumulation of saprotrophic fungi over the season may position these taxa to extract nutrients from the leaf tissue after it dies (Davis et al., 2023; Weatherhead et al., 2022), perhaps accelerating nutrient cycling by gaining priority over saprotrophic fungi in the soil that can colonize the leaf tissue only after it has entered the litter pool (Weatherhead et al., 2022).

The complete community was also much more variable over time, which included recruiting and losing taxa between survey dates. These stochastically present species are known as conditionally rare taxa and have been found to disproportionately contribute to temporal changes in the microbiome (Shade et al., 2014; Zuo et al., 2022). While we did not identify indicator species between the survey dates, there were distinct fungal lineages associated with early and late season communities in the phylogenetic neural networks, suggesting that, for some taxa, shared evolutionary history can help predict their role in seasonal assembly of the microbiome. *Cladosporidium* was highly predictive of early season communities and *Bulleribasidium* was most predictive of late season communities. Both taxa were identified within the core community, suggesting that they are persistently present but shifting in prevalence across the season. *Cladosporium* is an abundant genus in the soil and plant tissue that has been identified as both an endophyte (Clarke et al., 2006) and plant pathogen (Varga & Fischl, 2006) of tall fescue. Additional research has found *Bulleribasidium* to be a parasite of *Cladosporium* (Sampaio et al., 2002) suggesting parasitism could be the mechanism for the decline in *Cladosporium* and the rise in *Bulleribasidium* from the start to the end of the season. Combining our understanding of core and conditionally rare taxa with further analysis of trophic and transmission mode of predictive taxa could help us to identify the mechanisms leading recruitment and loss of taxa in the phyllosphere as plants grow throughout a season.

While fungal community diversity and dispersion were influenced by soil nutrient addition, we did not find any effect of soil nutrients on fungal trophic mode, or find any fungal taxa to be predictive of unfertilized or high fertilizer communities. Taxonomic richness of both the complete and core communities was significantly affected by nutrients and the interaction between nutrients and survey date. Richness of the low nutrient plots increased more slowly than richness of either the unfertilized or high nutrient plots, suggesting a non-linear effect of nutrients on the phyllosphere. Dispersion of the communities from the centroid was also influenced by nutrient addition with composition of the unfertilized plots experiencing increased dispersion compared to the low and high nutrient plots. Since these trends were observed for both the complete and core communities, it does not support conditionally rare species being responsible for the interaction with nutrients and instead suggests that both common and rare species of fungi are responding to soil nutrients.

Previous studies have found a consistent effect of nutrients on the soil and phyllosphere community at the site level (Ramirez et al., 2012; Seabloom et al., 2023). Our findings support this conclusion but also advise a more nuanced approach to future studies, since the response to nutrient addition was nonlinear. Community richness increased with low nutrient addition and community dispersion decreased with low and high nutrient addition. We applied a complete nutrient N-P-K fertilizer and thus cannot separate the effect of individual nutrients on the fungal community (Dordas, 2008). Additionally, we did not find an effect of nutrients on trophic mode of the foliar fungi. We found an accumulation of saprotrophic fungi throughout the season that was robust to nutrient addition.

When assessed in relationship to each other these results help to address the nexus of effects that seasonality and soil nutrients have on the phyllosphere microbiome. We found strong support for our first hypothesis that fungal diversity would increase early in the season as leaves recruit more taxa and identified the difference in the temporal variation of taxa richness between the complete and core communities. We also found support for our second hypothesis that a core community would establish early in the season and found that soil nutrients interacted with seasonality as the core community diverged throughout the season. Finally, trophic mode diversity decreased from early- to late-season communities and identified specific taxa that were predictive of early- and late-season communities, but not of nutrient addition. Further research into the seasonality of the phyllosphere at multiple sites and across multiple years could help us to better understand succession of fungal communities, how the microbiome reacts to and protects against seasonal epidemics of foliar pathogens, and how saprotrophic foliar and soil fungi interact as plant material joins the leaf litter (Davis et al., 2023; Debray et al., 2022; Trivedi et al., 2020; Weatherhead et al., 2022). Investigating the role that each of the core taxa and the predictive taxa play within the microbiome community could illuminate mechanisms by which the phyllosphere microbiome influences functioning of the leaf, in the context of biotic and abiotic stress (Custer et al., 2023; Howe et al., 2023; Shade & Handelsman, 2012).

## Supplementary Materials

**phyllosphere_microbiome_supplemental.docx**

**Figure S1.** Plot layout.

**Figure S2.** The proportion of total Bray-Curtis dissimilarity explained by the cumulative core community and by the complete community.

**Figure S3.** Distribution of sampling depth.

**Figure S4.** NMDS based on Bray-Curtis dissimilarity of fungal taxa by nutrient treatment.

**Table S1.** Linear mixed-effect model of Shannon diversity of plant species.

**Table S2.** PERMANOVA of Bray-Curtis dissimilarity of the plant community.

**Table S3.** ASV phylum assignments based on four classification methods.

**Table S4.** The 54 ASVs found in the core community, sorted by index number.

**Table S5.** Model of Shannon diversity indices of the complete community.

**Table S6.** Model of Shannon diversity indices of the core community.

**Table S7.** Model of taxa richness of the complete community.

**Table S8.** Model of taxa richness of the core community.

**Table S9.** Model of taxa evenness of the complete community.

**Table S10.** Model of taxa evenness of the core community.

**Table S11.** PERMANOVA of Bray-Curtis dissimilarity between the complete communities.

**Table S12.** Model of distance to centroid (dispersion) of Bray-Curtis dissimilarity of the complete communities.

**Table S13.** PERMANOVA of Bray-Curtis dissimilarity between the core communities.

**Table S14.** model of distance to centroid (dispersion) of Bray-Curtis dissimilarity of the core communities.

**Table S15.** Multivariate ANOVA with richness of each trophic mode in the complete community.

**Table S16.** Multivariate ANOVA with richness of each trophic mode in the core community.

**Figure S5.pdf**

**Figure S5.** Strength of predictive value of fungal lineages are determined through PopPhy-CNN.

**Table S17.xlsx**

**Table S17.** Feature scores for survey date and nutrients.

## Acknowledgements

This work was supported by the NSF-USDA joint program in Ecology and Evolution of Infectious Diseases (USDA-NIFA AFRI grant no. 2016-67013-25762 and NSF grant DEB-2308472). Development and ongoing enhancement of T-BAS was also supported by the Novo Nordisk Foundation (grant nos. NNF19SA0059360 [INTERACT] and NNF19SA0035476 [CCRP]) and by the U.S.–Israel Binational Agricultural Research and Development Fund (BARD) (Grant No. IS-5615-23R). We would like to thank E. Snyder and B. Stiver for help with sample collection, J. White for integrating PopPhy-CNN into T-BAS, and A. Hurlbert and S. McCoy for their written comments.

## Data Accessibility

Raw sequence reads, processed data, code, and related metadata are available on DataDryad (https://doi.org/10.5061/dryad.02v6wwqfd).

Benefits Generated: Benefits from this research accrue from the sharing of our data and results on public databases as described above.

## Author Contributions

E.T.G and C.E.M. conceived ideas and designed methodology; E.T.G. performed field collections and laboratory work; E.T.G. performed analyses and led writing of the manuscript.

I.C. performed additional analyses. All authors contributed critically to drafts and gave final

## Notes

### Competing Interest Statement

The authors have declared no competing interest.

### Summary of Updates

This version has been updated in response to reviewer comments. We have updated the introduction to highlight novelty and added additional analyses of fungal trophic mode (Figure 6). We also rearranged the results around key research questions.

## References

1. Argiroff, W. A., Carrell, A. A., Klingeman, D. M., Dove, N. C., Muchero, W., Veach, A. M., Wahl, T., Lebreux, S. J., Webb, A. B., Peyton, K., Schadt, C. W., & Cregger, M. A. (2024). Seasonality and longer-term development generate temporal dynamics in the Populus microbiome. mSystems, 9(3), e00886–23. 10.1128/msystems.00886-23

2. Arnillas, C. A., Borer, E. T., Seabloom, E. W., Alberti, J., Baez, S., Bakker, J. D., Boughton, E. H., Buckley, Y. M., Bugalho, M. N., Donohue, I., Dwyer, J., Firn, J., Gridzak, R., Hagenah, N., Hautier, Y., Helm, A., Jentsch, A., Knops, J. M. H., Komatsu, K. J., … Cadotte, M. W. (2021). Opposing community assembly patterns for dominant and nondominant plant species in herbaceous ecosystems globally. Ecology and Evolution, 11(24), 17744–17761. 10.1002/ece3.8266

3. Bates, D., Mächler, M., Bolker, B., & Walker, S. (2015). Fitting Linear Mixed-Effects Models Using lme4. Journal of Statistical Software, 67, 1–48. 10.18637/jss.v067.i01

4. Berg, M., & Koskella, B. (2018). Nutrient- and Dose-Dependent Microbiome-Mediated Protection against a Plant Pathogen. Current Biology, 28(15), 2487–2492.e3. 10.1016/j.cub.2018.05.085

5. Bolyen, E., Rideout, J. R., Dillon, M. R., Bokulich, N. A., Abnet, C. C., Al-Ghalith, G. A., Alexander, H., Alm, E. J., Arumugam, M., Asnicar, F., Bai, Y., Bisanz, J. E., Bittinger, K., Brejnrod, A., Brislawn, C. J., Brown, C. T., Callahan, B. J., Caraballo-Rodríguez, A. M., Chase, J., … Caporaso, J. G. (2019). Reproducible, interactive, scalable and extensible microbiome data science using QIIME 2. Nature Biotechnology, 37(8), Article 8. 10.1038/s41587-019-0209-9

6. Borer, E. T., Kinkel, L. L., May, G., & Seabloom, E. W. (2013). The world within: Quantifying the determinants and outcomes of a host’s microbiome. Basic and Applied Ecology, 14(7), 533–539. 10.1016/j.baae.2013.08.009

7. Boutin, S., & Laforest-Lapointe, I. (2025). Reproducing plant microbiome research reveals site and time as key drivers of apple tree phyllosphere bacterial communities. Scientific Reports, 15(1), 25620. 10.1038/s41598-025-10729-0

8. Busby, P. E., Peay, K. G., & Newcombe, G. (2016). Common foliar fungi of Populus trichocarpa modify Melampsora rust disease severity. The New Phytologist, 209(4), 1681–1692. 10.1111/nph.13742

9. Cáceres, M. D., & Legendre, P. (2009). Associations between species and groups of sites: Indices and statistical inference. Ecology, 90(12), 3566–3574. 10.1890/08-1823.1

10. Callahan, B. J., McMurdie, P. J., Rosen, M. J., Han, A. W., Johnson, A. J. A., & Holmes, S. P. (2016). DADA2: High-resolution sample inference from Illumina amplicon data. Nature Methods, 13(7), Article 7. 10.1038/nmeth.3869

11. Camacho, C., Coulouris, G., Avagyan, V., Ma, N., Papadopoulos, J., Bealer, K., & Madden, T. L. (2009). BLAST+: Architecture and applications. BMC Bioinformatics, 10, 421. 10.1186/1471-2105-10-421

12. Cantoran, A., Maillard, F., Baldrian, P., & Kennedy, P. G. (2023). Defining a core microbial necrobiome associated with decomposing fungal necromass. FEMS Microbiology Ecology, 99(9), fiad098. 10.1093/femsec/fiad098

13. Carbone, I., & White, J. B. (2025). T-BAS reference tree datasets [Data set]. T-BAS. 10.52750/634503.

14. Carbone, I., White, J. B., Miadlikowska, J., Arnold, A. E., Miller, M. A., Magain, N., U’Ren, J. M., & Lutzoni, F. (2019). T-BAS Version 2.1: Tree-Based Alignment Selector Toolkit for Evolutionary Placement of DNA Sequences and Viewing Alignments and Specimen Metadata on Curated and Custom Trees. Microbiology Resource Announcements, 8(29), e00328–19. 10.1128/MRA.00328-19

15. Clarke, B. B., White, J. F., Hurley, R. H., Torres, M. S., Sun, S., & Huff, D. R. (2006). Endophyte-Mediated Suppression of Dollar Spot Disease in Fine Fescues. Plant Disease, 90(8), 994–998. 10.1094/PD-90-0994

16. Copeland, J. K., Yuan, L., Layeghifard, M., Wang, P. W., & Guttman, D. S. (2015). Seasonal Community Succession of the Phyllosphere Microbiome. Molecular Plant-Microbe Interactions®, 28(3), 274–285. 10.1094/MPMI-10-14-0331-FI

17. Custer, G. F., Gans, M., van Diepen, L. T. A., Dini-Andreote, F., & Buerkle, C. A. (2023). Comparative Analysis of Core Microbiome Assignments: Implications for Ecological Synthesis. mSystems, 8(1), e01066–22. 10.1128/msystems.01066-22

18. Dale, J. C. M., & Newman, J. A. (2022). A First Draft of the Core Fungal Microbiome of Schedonorus arundinaceus with and without Its Fungal Mutualist Epichloë coenophiala. Journal of Fungi, 8(10), Article 10. 10.3390/jof8101026

19. Davis, E. L., Weatherhead, E., & Koide, R. T. (2023). The potential saprotrophic capacity of foliar endophytic fungi from *Quercus gambelii*. Fungal Ecology, 62, 101221. 10.1016/j.funeco.2022.101221

20. Debray, R., Herbert, R. A., Jaffe, A. L., Crits-Christoph, A., Power, M. E., & Koskella, B. (2022). Priority effects in microbiome assembly. Nature Reviews Microbiology, 20(2), Article 2. 10.1038/s41579-021-00604-w

21. Dordas, C. (2008). Role of nutrients in controlling plant diseases in sustainable agriculture. A review. Agronomy for Sustainable Development, 28, 33–46.

22. Fang, K., Zhou, J., Chen, L., Li, Y.-X., Yang, A.-L., Dong, X.-F., & Zhang, H.-B. (2021). Virulence and community dynamics of fungal species with vertical and horizontal transmission on a plant with multiple infections. PLOS Pathogens, 17(7), e1009769. 10.1371/journal.ppat.1009769

23. Firn, J., McGree, J. M., Harvey, E., Flores-Moreno, H., Schütz, M., Buckley, Y. M., Borer, E. T., Seabloom, E. W., La Pierre, K. J., MacDougall, A. M., Prober, S. M., Stevens, C. J., Sullivan, L. L., Porter, E., Ladouceur, E., Allen, C., Moromizato, K. H., Morgan, J. W., Harpole, W. S., … Risch, A. C. (2019). Leaf nutrients, not specific leaf area, are consistent indicators of elevated nutrient inputs. Nature Ecology & Evolution, 3(3), Article 3. 10.1038/s41559-018-0790-1

24. Fürnkranz, M., Wanek, W., Richter, A., Abell, G., Rasche, F., & Sessitsch, A. (2008). Nitrogen fixation by phyllosphere bacteria associated with higher plants and their colonizing epiphytes of a tropical lowland rainforest of Costa Rica. The ISME Journal, 2(5), Article 5. 10.1038/ismej.2008.14

25. Gao, C., Montoya, L., Xu, L., Madera, M., Hollingsworth, J., Purdom, E., Singan, V., Vogel, J., Hutmacher, R. B., Dahlberg, J. A., Coleman-Derr, D., Lemaux, P. G., & Taylor, J. W. (2020). Fungal community assembly in drought-stressed sorghum shows stochasticity, selection, and universal ecological dynamics. Nature Communications, 11(1), 34. 10.1038/s41467-019-13913-9

26. Gao, F.-L., Che, X.-X., Yu, F.-H., & Li, J.-M. (2019). Cascading effects of nitrogen, rhizobia and parasitism via a host plant. Flora, 251, 62–67. 10.1016/j.flora.2018.12.007

27. Geyer, J. K., Grunberg, R. L., Wang, J., & Mitchell, C. E. (2024). Leaf age structures phyllosphere microbial communities in the field and greenhouse. Frontiers in Microbiology, 15. 10.3389/fmicb.2024.1429166

28. Grady, K. L., Sorensen, J. W., Stopnisek, N., Guittar, J., & Shade, A. (2019). Assembly and seasonality of core phyllosphere microbiota on perennial biofuel crops. Nature Communications, 10(1), 4135. 10.1038/s41467-019-11974-4

29. Grunberg, R. L., Halliday, F. W., Heckman, R. W., Joyner, B. N., O’Keeffe, K. R., & Mitchell, C. E. (2023). Disease decreases variation in host community structure in an old-field grassland. PLOS ONE, 18(10), e0293495. 10.1371/journal.pone.0293495

30. Halliday, F. W., Heckman, R. W., Wilfahrt, P. A., & Mitchell, C. E. (2019). Past is prologue: Host community assembly and the risk of infectious disease over time. Ecology Letters, 22(1), 138–148. 10.1111/ele.13176

31. Heckman, R. W., Halliday, F. W., & Mitchell, C. E. (2019). A growth–defense trade-off is general across native and exotic grasses. Oecologia, 191(3), 609–620. 10.1007/s00442-019-04507-9

32. Heckman, R. W., Wright, J. P., & Mitchell, C. E. (2016). Joint effects of nutrient addition and enemy exclusion on exotic plant success. Ecology, 97(12), 3337–3345. 10.1002/ecy.1585

33. Hernandez-Agreda, A., Gates, R. D., & Ainsworth, T. D. (2017). Defining the Core Microbiome in Corals’ Microbial Soup. Trends in Microbiology, 25(2), 125–140. 10.1016/j.tim.2016.11.003

34. Hodgson, S., Cates, C., Hodgson, J., Morley, N. J., Sutton, B. C., & Gange, A. C. (2014). Vertical transmission of fungal endophytes is widespread in forbs. Ecology and Evolution, 4(8), 1199–1208. 10.1002/ece3.953

35. Howe, A., Stopnisek, N., Dooley, S. K., Yang, F., Grady, K. L., & Shade, A. (2023). Seasonal activities of the phyllosphere microbiome of perennial crops. Nature Communications, 14, 1039. 10.1038/s41467-023-36515-y

36. Humphrey, P. T., Nguyen, T. T., Villalobos, M. M., & Whiteman, N. K. (2014). Diversity and abundance of phyllosphere bacteria are linked to insect herbivory. Molecular Ecology, 23(6), 1497–1515. 10.1111/mec.12657

37. James, T. Y., Kauff, F., Schoch, C. L., Matheny, P. B., Hofstetter, V., Cox, C. J., Celio, G., Gueidan, C., Fraker, E., Miadlikowska, J., Lumbsch, H. T., Rauhut, A., Reeb, V., Arnold, A. E., Amtoft, A., Stajich, J. E., Hosaka, K., Sung, G.-H., Johnson, D., … Vilgalys, R. (2006). Reconstructing the early evolution of Fungi using a six-gene phylogeny. Nature, 443(7113), 818–822. 10.1038/nature05110

38. Keller, A. B., Walter, C. A., Blumenthal, D. M., Borer, E. T., Collins, S. L., DeLancey, L. C., Fay, P. A., Hofmockel, K. S., Knops, J. M. H., Leakey, A. D. B., Mayes, M. A., Seabloom, E. W., & Hobbie, S. E. (2023). Stronger fertilization effects on aboveground versus belowground plant properties across nine U.S. grasslands. Ecology, 104(2), e3891. 10.1002/ecy.3891

39. Kong, H. G., Song, G. C., & Ryu, C.-M. (2019). Inheritance of seed and rhizosphere microbial communities through plant–soil feedback and soil memory. Environmental Microbiology Reports, 11(4), 479–486. 10.1111/1758-2229.12760

40. Lacroix, C., Seabloom, E. W., & Borer, E. T. (2014). Environmental nutrient supply alters prevalence and weakens competitive interactions among coinfecting viruses. New Phytologist, 204(2), 424–433. 10.1111/nph.12909

41. Liu, X., Lyu, S., Sun, D., Bradshaw, C. J. A., & Zhou, S. (2017). Species decline under nitrogen fertilization increases community-level competence of fungal diseases. Proceedings of the Royal Society B: Biological Sciences, 284(1847), 20162621. 10.1098/rspb.2016.2621

42. Love, M. I., Huber, W., & Anders, S. (2014). Moderated estimation of fold change and dispersion for RNA-seq data with DESeq2. Genome Biology, 15(12), 550. 10.1186/s13059-014-0550-8

43. Lumibao, C. Y., Borer, E. T., Condon, B., Kinkel, L., May, G., & Seabloom, E. W. (2019). Site specific responses of foliar fungal microbiomes to nutrient addition and herbivory at different spatial scales. Ecology and Evolution, 9(21), 12231–12244. 10.1002/ece3.5711

44. Lundberg, D. S., Lebeis, S. L., Paredes, S. H., Yourstone, S., Gehring, J., Malfatti, S., Tremblay, J., Engelbrektson, A., Kunin, V., del Rio, T. G., Edgar, R. C., Eickhorst, T., Ley, R. E., Hugenholtz, P., Tringe, S. G., & Dangl, J. L. (2012). Defining the core Arabidopsis thaliana root microbiome. Nature, 488(7409), 86–90. 10.1038/nature11237

45. Marañón-Jiménez, S., Radujković, D., Verbruggen, E., Grau, O., Cuntz, M., Peñuelas, J., Richter, A., Schrumpf, M., & Rebmann, C. (2021). Shifts in the Abundances of Saprotrophic and Ectomycorrhizal Fungi With Altered Leaf Litter Inputs. Frontiers in Plant Science, 12. 10.3389/fpls.2021.682142

46. Mitchell, C. E., Tilman, D., & Groth, J. V. (2002). Effects of Grassland Plant Species Diversity, Abundance, and Composition on Foliar Fungal Disease. Ecology, 83(6), 1713–1726. 10.1890/0012-9658(2002)083%255B1713:EOGPSD%255D2.0.CO;2

47. Nguyen, N. H., Song, Z., Bates, S. T., Branco, S., Tedersoo, L., Menke, J., Schilling, J. S., & Kennedy, P. G. (2016). FUNGuild: An open annotation tool for parsing fungal community datasets by ecological guild. Fungal Ecology, 20, 241–248. 10.1016/j.funeco.2015.06.006

48. Oksanen, J., Simpson, G. L., Blanchet, F. G., Kindt, R., Legendre, P., Minchin, P. R., O’Hara, R. B., Solymos, P., Stevens, M. H. H., Szoecs, E., Wagner, H., Barbour, M., Bedward, M., Bolker, B., Borcard, D., Carvalho, G., Chirico, M., Caceres, M. D., Durand, S., … Weedon, J. (2022). *vegan: Community Ecology Package* (Version 2.6-4) [Computer software]. https://cran.r-project.org/web/packages/vegan/index.html

49. Oono, R., Black, D., Slessarev, E., Sickler, B., Strom, A., & Apigo, A. (2020). Species diversity of fungal endophytes across a stress gradient for plants. New Phytologist, 228(1), 210–225. 10.1111/nph.16709

50. Otten, W., Hall, D., Harris, K., Ritz, K., Young, I. M., & Gilligan, C. A. (2001). Soil physics, fungal epidemiology and the spread of *Rhizoctonia solani*. New Phytologist, 151(2), 459–468. 10.1046/j.0028-646x.2001.00190.x

51. Põlme, S., Abarenkov, K., Henrik Nilsson, R., Lindahl, B. D., Clemmensen, K. E., Kauserud, H., Nguyen, N., Kjøller, R., Bates, S. T., Baldrian, P., Frøslev, T. G., Adojaan, K., Vizzini, A., Suija, A., Pfister, D., Baral, H.-O., Järv, H., Madrid, H., Nordén, J., … Tedersoo, L. (2020). FungalTraits: A user-friendly traits database of fungi and fungus-like stramenopiles. Fungal Diversity, 105(1), 1–16. 10.1007/s13225-020-00466-2

52. Ramirez, K. S., Craine, J. M., & Fierer, N. (2012). Consistent effects of nitrogen amendments on soil microbial communities and processes across biomes. Global Change Biology, 18(6), 1918–1927. 10.1111/j.1365-2486.2012.02639.x

53. Reiman, D., Metwally, A. A., Sun, J., & Dai, Y. (2020). PopPhy-CNN: A Phylogenetic Tree Embedded Architecture for Convolutional Neural Networks to Predict Host Phenotype From Metagenomic Data. IEEE Journal of Biomedical and Health Informatics, 24(10), 2993–3001. 10.1109/JBHI.2020.2993761

54. Ritpitakphong, U., Falquet, L., Vimoltust, A., Berger, A., Métraux, J.-P., & L’Haridon, F. (2016). The microbiome of the leaf surface of Arabidopsis protects against a fungal pathogen. New Phytologist, 210(3), 1033–1043. 10.1111/nph.13808

55. Sampaio, J. P., Weiß, M., Gadanho, M., & Bauer, R. (2002). New taxa in the Tremellales: Bulleribasidium oberjochense gen. Et sp. Nov., Papiliotrema bandonii ge. Mycologia, 94(5), 873–887.

56. Seabloom, E. W., Caldeira, M. C., Davies, K. F., Kinkel, L., Knops, J. M. H., Komatsu, K. J., MacDougall, A. S., May, G., Millican, M., Moore, J. L., Perez, L. I., Porath-Krause, A. J., Power, S. A., Prober, S. M., Risch, A. C., Stevens, C., & Borer, E. T. (2023). Globally consistent response of plant microbiome diversity across hosts and continents to soil nutrients and herbivores. Nature Communications, 14(1), Article 1. 10.1038/s41467-023-39179-w

57. Shade, A., & Handelsman, J. (2012). Beyond the Venn diagram: The hunt for a core microbiome. Environmental Microbiology, 14(1), 4–12. 10.1111/j.1462-2920.2011.02585.x

58. Shade, A., Jones, S. E., Caporaso, J. G., Handelsman, J., Knight, R., Fierer, N., & Gilbert, J. A. (2014). Conditionally Rare Taxa Disproportionately Contribute to Temporal Changes in Microbial Diversity. mBio, 5(4), 10.1128/mbio.01371-14. 10.1128/mbio.01371-14

59. Shade, A., & Stopnisek, N. (2019). Abundance-occupancy distributions to prioritize plant core microbiome membership. Current Opinion in Microbiology, 49, 50–58. 10.1016/j.mib.2019.09.008

60. Shannon, P., Markiel, A., Ozier, O., Baliga, N. S., Wang, J. T., Ramage, D., Amin, N., Schwikowski, B., & Ideker, T. (2003). Cytoscape: A software environment for integrated models of biomolecular interaction networks. Genome Research, 13(11), 2498–2504. 10.1101/gr.1239303

61. Shillcock, G., Úbeda, F., & Wild, G. (2023). Vertical transmission does not always lead to benign pathogen–host associations. Evolution Letters, 7(5), 305–314. 10.1093/evlett/qrad028

62. Smith, K. F., Acevedo-Whitehouse, K., & Pedersen, A. B. (2009). The role of infectious diseases in biological conservation. Animal Conservation, 12(1), 1–12. 10.1111/j.1469-1795.2008.00228.x

63. Steffen, W., Richardson, K., Rockström, J., & Sarah E. Cornell. (2015). Planetary boundaries: Guiding human development on a changing planet. 347(6223), 736. 10.1126/science.1259855

64. Tellez, P. H., Arnold, A. E., Leo, A. B., Kitajima, K., & Van Bael, S. A. (2022). Traits along the leaf economics spectrum are associated with communities of foliar endophytic symbionts. Frontiers in Microbiology, 13. 10.3389/fmicb.2022.927780

65. Trivedi, P., Leach, J. E., Tringe, S. G., Sa, T., & Singh, B. K. (2020). Plant–microbiome interactions: From community assembly to plant health. Nature Reviews Microbiology, 18(11), Article 11. 10.1038/s41579-020-0412-1

66. Umaña, M. N., Zhang, C., Cao, M., Lin, L., & Swenson, N. G. (2017). A core-transient framework for trait-based community ecology: An example from a tropical tree seedling community. Ecology Letters, 20(5), 619–628. 10.1111/ele.12760

67. Varga, Z., & Fischl, G. (2006). Fungal infection of cultivated grass seeds. Cereal Research Communications, 34(4), 1291–1297. 10.1556/CRC.34.2006.4.271

68. Veresoglou, S. D., Barto, E. K., Menexes, G., & Rillig, M. C. (2013). Fertilization affects severity of disease caused by fungal plant pathogens. Plant Pathology, 62(5), 961–969. 10.1111/ppa.12014

69. Wagner, M. R., Busby, P. E., & Balint Kurti, P. (2020). Analysis of leaf microbiome composition of near-isogenic maize lines differing in broad-spectrum disease resistance. New Phytologist, 225(5), 2152–2165. 10.1111/nph.16284

70. Wang, Q., & Cole, J. R. (2024). Updated RDP taxonomy and RDP Classifier for more accurate taxonomic classification. Microbiology Resource Announcements, 13(4), e01063–23. 10.1128/mra.01063-23

71. Wang, S., Tan, Y., Luo, Q., Fang, X., Zhu, H., Li, S., Zhou, Y., & Zhu, T. (2025). Temporal dynamics of walnut phyllosphere microbiota under synergistic pathogen exposure and environmental perturbation. Frontiers in Microbiology, 16, 1551476. 10.3389/fmicb.2025.1551476

72. Washburne, A. D., Morton, J. T., Sanders, J., McDonald, D., Zhu, Q., Oliverio, A. M., & Knight, R. (2018). Methods for phylogenetic analysis of microbiome data. Nature Microbiology, 3(6), 652–661. 10.1038/s41564-018-0156-0

73. Weatherhead, E., Davis, E. L., & Koide, R. T. (2022). Many foliar endophytic fungi of Quercus gambelii are capable of psychrotolerant saprotrophic growth. PloS One, 17(10), e0275845. 10.1371/journal.pone.0275845

74. Xiao, J., Chen, L., Johnson, S., Yu, Y., Zhang, X., & Chen, J. (2018). Predictive Modeling of Microbiome Data Using a Phylogeny-Regularized Generalized Linear Mixed Model. Frontiers in Microbiology, 9, 1391. 10.3389/fmicb.2018.01391

75. Zuo, Y., Zeng, R., Tian, C., Wang, J., & Qu, W. (2022). The importance of conditionally rare taxa for the assembly and interaction of fungal communities in mangrove sediments. Applied Microbiology and Biotechnology, 106(9), 3787–3798. 10.1007/s00253-022-11949-4

