## Supplementary material for "Seasonal assembly of the phyllosphere fungal microbiome of a perennial grass is robust to nutrient addition": 05-05-25 microbiome supplemental.pdf

**Supplemental Figures**

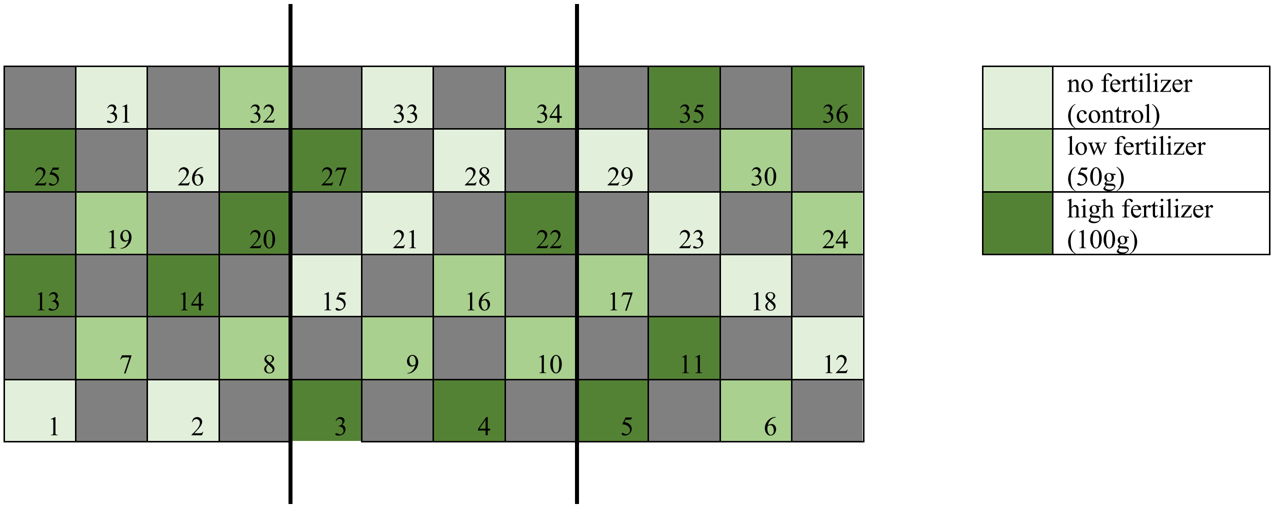


**Figure S1.** Plot layout. Plots were arranged in a checkerboard pattern and colors represent nutrient addition treatments. Treatments were randomly assigned within three spatial blocks. Blocks occur from left to right and are separated by two longer vertical lines.

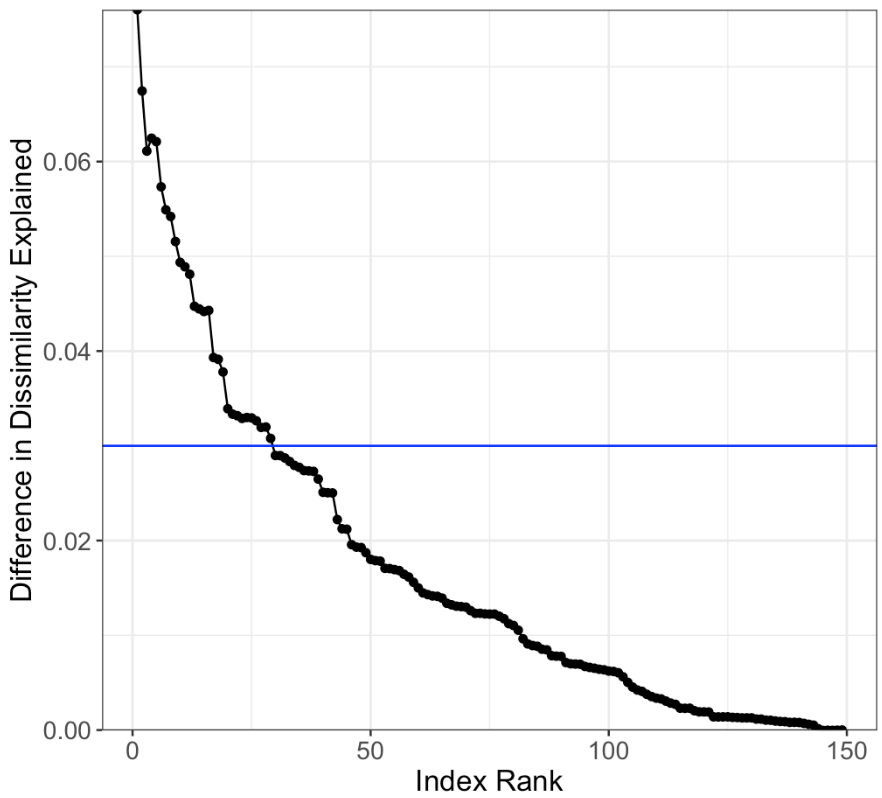


**Figure S2.** The difference between the proportion of total Bray-Curtis dissimilarity explained by the cumulative core community and the proportion explained by the complete community. The blue horizontal line indicates the cut-off for inclusion in the core community: where including the next-ranked ASVs would have added less than 0.03 to the proportion explained, which was after ASV index rank 28. Adding additional taxa to the core community failed to explain additional variation in Bray-Curtis dissimilarly.

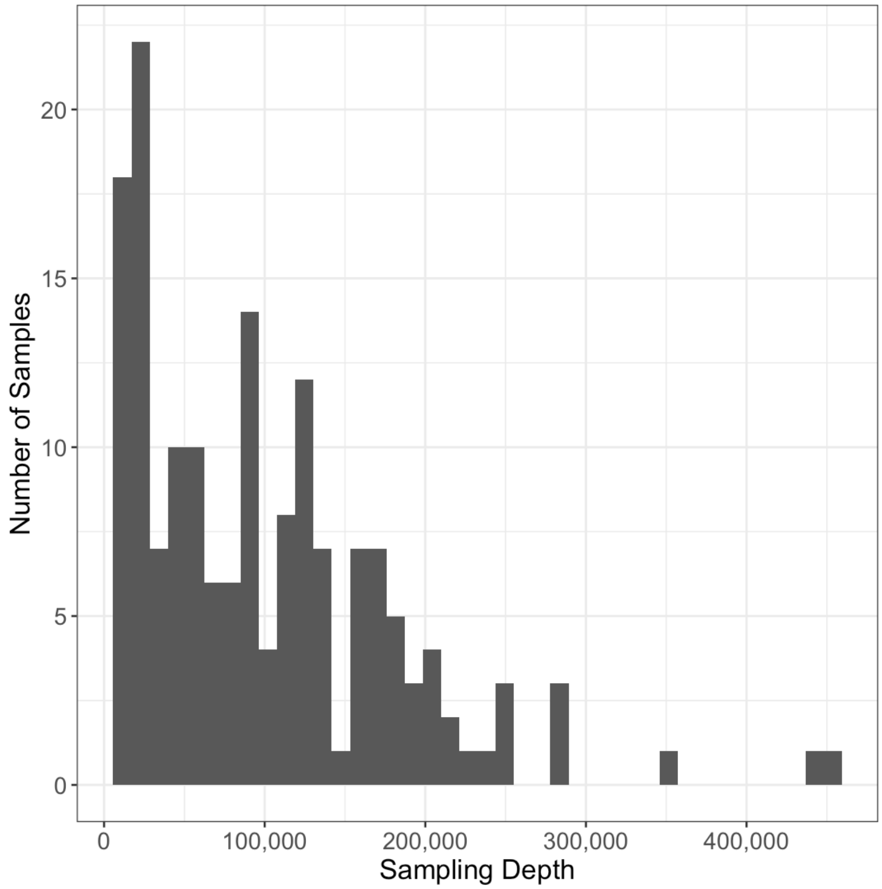


**Figure S3.** Distribution of sampling depth (number of reads) for all samples remaining after filtering.

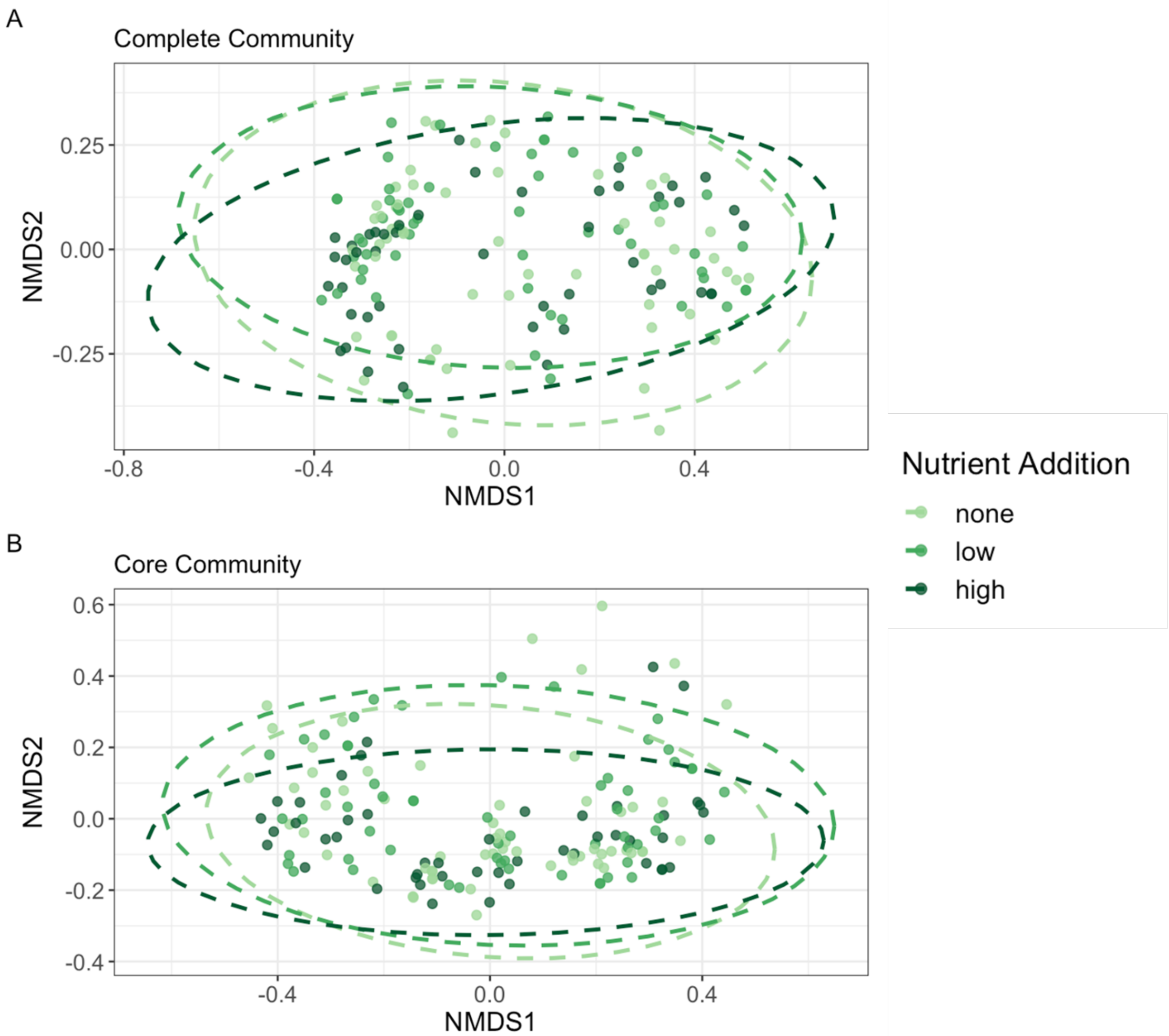


**Figure S4.** NMDS based on Bray-Curtis dissimilarity of fungal taxa for the complete (A) and the core (B) fungal taxa in each plot, grouped by survey date. Each point represents a plot and color represents the nutrient treatment. 95% confidence ellipses are shown for each nutrient group.

**Figure S5.** Strength of predictive value of fungal lineages are determined through PopPhy-CNN for early and late season communities (A; n=117) and low and high nutrient addition communities (B; n=72). The tree was drawn to reflect the lineage hierarchy, starting with taxa (ASVs) at the tips, then following nodes inward leads through genus, family, order, class, and phylum (Table S1). Vibrancy of blue edges illustrates strength of predictive value for early season and low nutrient addition communities and vibrancy of orange edges illustrates strength of predictive value for late season and high nutrient communities.

### Supplemental Tables

**Table S1.** ASV phylum assignments based on four classification methods.

| Taxon | UNITE | T-BAS | RDP Classifier | NCBI |
|---|---|---|---|---|
| Ascomycota | 2102 | 2131 | 2045 | 1963 |
| Basidiomycota | 968 | 969 | 1008 | 979 |
| Mucoromycota | 0 | 23 | 0 | 11 |
| Zoopagomycota | 0 | 9 | 0 | 0 |
| Chytridiomycota | 6 | 6 | 6 | 7 |
| Dikarya incertae sedis | 0 | 1 | 0 | 0 |
| Rozellomycota | 4 | 0 | 0 | 0 |
| Mortierellomycota | 6 | 0 | 0 | 0 |
| Glomeromycota | 13 | 0 | 72 | 248 |
| Zygomycota | 0 | 0 | 14 | 2 |
| unidentified | 40 | 0 | 110 | 0 |

**Table S2.** The 54 ASVs found in the core community, sorted by index number. Colors correspond to the taxonomic order in which they were placed (orange = Pleosporales, pink = Hypocreales, green = Tremellales, blue = Capniodiales, purple = all orders with only one placement, gray = taxonomic order could not be determined).

| Index | ASV | Phylum | Class | Order | Family | Genus |
|---|---|---|---|---|---|---|
| 1 | 1 | Ascomycota | Dothideomycetes | Pleosporales | Didymellaceae | Neoascochyta |
| 1 | 10 | Ascomycota | Dothideomycetes | Pleosporales | Pleosporaceae | Alternaria |
| 1 | 11 | Ascomycota | Sordariomycetes | Hypocreales | Hypocreales_fam_Incertae_sedis | Sarocladium |
| 1 | 2 | Ascomycota | Dothideomycetes | Pleosporales | Didymellaceae | Ascochyta |
| 1 | 24 | Ascomycota | Dothideomycetes | Pleosporales | Didymellaceae | Neoascochyta |
| 1 | 26 | Basidiomycota | Tremellomycetes | Tremellales | Bulleraceae | Bullera |
| 1 | 3 | Ascomycota | Dothideomycetes | Capnodiales | Cladosporiaceae | Cladosporium |

| | | | | | | |
|---|---|---|---|---|---|---|
| 1 | 4 | Basidiomycota | Tremellomycetes | Tremellales | Bulleribasidiaceae | Vishniacozyma |
| 1 | 5 | Ascomycota | Dothideomycetes | Pleosporales | Didymellaceae | Epicoccum |
| 1 | 6 | Ascomycota | Dothideomycetes | Capnodiales | Mycosphaerellaceae | Zymoseptoria |
| 1 | 7 | Ascomycota | Dothideomycetes | Pleosporales | Cucurbitariaceae | Pyrenochaetopsis |
| 1 | 8 | Ascomycota | Dothideomycetes | Pleosporales | Cucurbitariaceae | Pyrenochaetopsis |
| 2 | 12 | Basidiomycota | Tremellomycetes | Tremellales | Bulleribasidiaceae | Vishniacozyma |
| 2 | 14 | Ascomycota | Dothideomycetes | Capnodiales | Mycosphaerellaceae | Mycosphaerella |
| 2 | 15 | Ascomycota | Dothideomycetes | Capnodiales | Dissoconiaceae | Dissoconium |
| 2 | 16 | Ascomycota | Dothideomycetes | Pleosporales | Didymellaceae | Neoascochyta |
| 2 | 19 | Ascomycota | Dothideomycetes | Capnodiales | Mycosphaerellaceae | Ramularia |
| 2 | 30 | Ascomycota | Dothideomycetes | Capnodiales | Neodevriesiaceae | Neodevriesia |
| 2 | 38 | Basidiomycota | Microbotryomycetes | Sporidiobolales | Sporidiobolaceae | Sporobolomyces |
| 2 | 61 | Basidiomycota | Tremellomycetes | Tremellales | Bulleribasidiaceae | Vishniacozyma |
| 3 | 23 | Ascomycota | Dothideomycetes | Pleosporales | Cucurbitariaceae | Pyrenochaetopsis |
| 3 | 27 | Ascomycota | Dothideomycetes | Pleosporales | Didymellaceae | Ascochyta |
| 3 | 48 | Basidiomycota | Tremellomycetes | Tremellales | Tremellaceae | Cryptococcus |
| 4 | 13 | Ascomycota | Dothideomycetes | Pleosporales | Didymellaceae | Epicoccum |
| 5 | 25 | Ascomycota | Dothideomycetes | Pleosporales | Phaeosphaeriaceae | Phaeosphaeria |
| 6 | 9 | Ascomycota | Sordariomycetes | Hypocreales | Nectriaceae | Gibberella |
| 7 | 22 | Basidiomycota | Tremellomycetes | Cystofilobasidiales | Cystofilobasidiaceae | Cystofilobasidium |
| 8 | 28 | Ascomycota | Dothideomycetes | Pleosporales | Didymellaceae | NA |
| 9 | 33 | Ascomycota | Sordariomycetes | Hypocreales | Hypocreales_fam_Incertae_sedis | Acremonium |
| 10 | 35 | Ascomycota | Dothideomycetes | Capnodiales | Teratosphaeriaceae | Apenidiella |
| 11 | 54 | Ascomycota | Dothideomycetes | Pleosporales | NA | NA |
| 12 | 39 | Ascomycota | Sordariomycetes | Hypocreales | Ophiocordycipitaceae | Hirsutella |
| 13 | 21 | Ascomycota | Sordariomycetes | Hypocreales | Nectriaceae | Fusarium |
| 14 | 47 | Ascomycota | Dothideomycetes | Capnodiales | Mycosphaerellaceae | Ramularia |
| 15 | 57 | Basidiomycota | Tremellomycetes | Tremellales | NA | NA |
| 16 | 32 | Ascomycota | Dothideomycetes | Pleosporales | Cucurbitariaceae | Pyrenochaetopsis |
| 16 | 53 | Basidiomycota | Tremellomycetes | Tremellales | Bulleribasidiaceae | Hannaella |
| 17 | 20 | Ascomycota | Dothideomycetes | Capnodiales | Mycosphaerellaceae | Cercospora |
| 18 | 58 | Ascomycota | Dothideomycetes | Capnodiales | Neodevriesiaceae | Neodevriesia |
| 18 | 69 | Ascomycota | Dothideomycetes | Pleosporales | NA | NA |
| 19 | 36 | Ascomycota | Sordariomycetes | Hypocreales | Stachybotryaceae | Myrothecium |

| | | | | | | |
|---|---|---|---|---|---|---|
| 19 | 62 | Ascomycota | Sordariomycetes | Glomerellales | Plectosphaerellaceae | Plectosphaerella |
| 20 | 17 | Ascomycota | Dothideomycetes | Pleosporales | Cucurbitariaceae | Pyrenochaetopsis |
| 21 | 45 | Ascomycota | Dothideomycetes | Pleosporales | NA | NA |
| 22 | 79 | Ascomycota | NA | NA | NA | NA |
| 22 | 84 | Basidiomycota | Cystobasidiomycetes | Erythrobasidiales | Erythrobasidiaceae | Bannoa |
| 23 | 46 | Ascomycota | Dothideomycetes | Capnodiales | Cladosporiaceae | Cladosporium |
| 24 | 42 | Ascomycota | Dothideomycetes | Pleosporales | Didymellaceae | Ascochyta |
| 24 | 49 | Ascomycota | Dothideomycetes | Pleosporales | NA | NA |
| 25 | 63 | Basidiomycota | Cystobasidiomycetes | NA | NA | NA |
| 26 | 59 | Ascomycota | Dothideomycetes | Pleosporales | Periconiaceae | Periconia |
| 26 | 31 | Ascomycota | Dothideomycetes | Capnodiales | Dissoconiaceae | Dissoconium |
| 27 | 177 | Basidiomycota | Agaricostilbomycetes | Agaricostilbales | Chionosphaeraceae | NA |
| 28 | 44 | Ascomycota | Dothideomycetes | Acrospermales | Acrospermales_fam_Incertae_sedis | Phaeodactylium |

**Table S3.** Quadratic mixed effect model of log transformed Shannon diversity indices of the complete community with sampling depth as a fixed effect and plot nested in spatial block as random intercepts and an ANOVA posthoc test (marginal $R^2 = 0.66$, conditional $R^2 = 0.81$). = log(Shannon diversity index) ~ Sampling Depth + (Date + Date$^2$) * Nutrients + (1 | Block / Plot) – ANOVA

| **Variable** | **df** | **Sum Sq** | **Mean Sq** | **F statistic** | **p-value** |
|---|---|---|---|---|---|
| Sampling Depth | 1 | 0.0501 | 0.0501 | 8.1825 | 0.0048 |
| Date | 1 | 0.7797 | 0.7797 | 127.2523 | <0.001 |
| Date$^2$ | 1 | 0.7277 | 0.7277 | 118.7589 | <0.001 |
| Nutrients | 2 | 0.0189 | 0.0095 | 1.5459 | 0.3927 |
| Date * Nutrients | 2 | 0.0065 | 0.0032 | 0.5275 | 0.5911 |
| Date$^2$ * Nutrients | 2 | 0.0189 | 0.0095 | 1.5467 | 0.3926 |

**Table S4.** Quadratic mixed effect model of log transformed Shannon diversity indices of the core community with sampling depth as a fixed effect and plot nested in spatial block as random intercepts and an ANOVA posthoc test (marginal $R^2 = 0.70$, conditional $R^2 = 0.73$).

= log(Shannon diversity index of core) ~ Sampling Depth + (Date + Date$^2$) * Nutrients + (1 | Block / Plot) – ANOVA

| **Variable** | **df** | **Sum Sq** | **Mean Sq** | **F statistic** | **p-value** |
|---|---|---|---|---|---|
| Sampling Depth | 1 | 0.0530 | 0.0530 | 6.9329 | 0.0325 |
| Date | 1 | 0.5833 | 0.5833 | 79.2349 | <0.001 |
| Date$^2$ | 1 | 1.0039 | 1.0039 | 131.2138 | <0.001 |
| Nutrients | 2 | 0.0207 | 0.0104 | 1.3549 | 0.4247 |
| Date * Nutrients | 2 | 0.0123 | 0.0062 | 0.8034 | 0.4639 |
| Date$^2$ * Nutrients | 2 | 0.0207 | 0.0104 | 1.3533 | 0.4249 |

**Table S5.** Linear mixed effect model of log transformed taxa richness of the complete community with sampling depth as a fixed effect and plot nested in spatial block as random intercepts and an ANOVA posthoc test (marginal $R^2 = 0.68$, conditional $R^2 = 0.76$).

= log(Richness) ~ Sampling Depth + Date * Nutrients + (1 | Block / Plot) – ANOVA

| **Variable** | **df** | **Sum Sq** | **Mean Sq** | **F statistic** | **p-value** |
|---|---|---|---|---|---|
| Sampling Depth | 1 | 0.8648 | 0.8648 | 137.60 | <0.001 |
| Date | 1 | 2.4230 | 2.4230 | 379.92 | <0.001 |
| Nutrients | 2 | 2.2019 | 1.1010 | 345.25 | <0.001 |
| Date * Nutrients | 2 | 2.2002 | 1.1001 | 172.49 | <0.001 |

**Table S6.** Quadratic mixed effect model of log transformed taxa richness of the core community with sampling depth as random intersects and plot nested in spatial block as random intercepts and an ANOVA posthoc test (marginal $R^2 = 0.63$, conditional $R^2 = 0.68$).

= log(Richness of Core) ~ Sampling Depth + (Date + Date$^2$) * Nutrients + (1 | Block / Plot) – ANOVA

| Nutrients | df | Sum Sq | Mean Sq | F statistic | p-value |
|---|---|---|---|---|---|
| Sampling Depth | 1 | 0.0295 | 0.0295 | 37.3663 | <0.001 |
| Date | 1 | 0.0206 | 0.0206 | 26.0879 | <0.001 |
| Date$^2$ | 1 | 0.0848 | 0.0848 | 107.5711 | <0.001 |
| Nutrients | 2 | 0.0001 | 0.0001 | 0.0164 | 0.9839 |
| Date * Nutrients | 2 | 0.0075 | 0.0037 | 4.7434 | 0.0102 |
| Date$^2$ * Nutrients | 2 | 0.0001 | 0.0001 | 0.0166 | 0.9837 |

**Table S7.** Quadratic mixed effect model of log transformed taxa evenness of the complete community with sampling depth as a fixed effect and plot nested in spatial block as random intercepts and an ANOVA posthoc test (marginal $R^2 = 0.72$, conditional $R^2 = 0.73$).

= log(Evenness) ~ Sampling Depth + (Date + Date$^2$) * Nutrients + (1 | Block / Plot) – ANOVA

| Nutrients | df | Sum Sq | Mean Sq | F statistic | p-value |
|---|---|---|---|---|---|
| Sampling Depth | 1 | 0.1799 | 0.1799 | 33.1810 | <0.001 |
| Date | 1 | 0.4657 | 0.4657 | 85.8972 | 0.0672 |
| Date$^2$ | 1 | 0.5738 | 0.5738 | 105.8342 | <0.001 |
| Nutrients | 2 | 0.0123 | 0.0062 | 1.1390 | 0.4675 |
| Date * Nutrients | 2 | 0.0036 | 0.0018 | 0.3352 | 0.7157 |
| Date$^2$ * Nutrients | 2 | 0.0123 | 0.0062 | 1.1374 | 0.4678 |

**Table S8.** Quadratic mixed effect model of log transformed taxa evenness of the core community with sampling depth as random intersects and plot nested in spatial block as random intercepts and an ANOVA posthoc test (marginal $R^2 = 0.69$, conditional $R^2 = 0.70$).

= log(Evenness of Core) ~ Sampling Depth + (Date + Date$^2$) * Nutrients + (1 | Block / Plot) – ANOVA

| **Variable** | **df** | **Sum Sq** | **Mean Sq** | **F statistic** | **p-value** |
|---|---|---|---|---|---|
| Sampling Depth | 1 | 0.0742 | 0.0742 | 10.1305 | <0.001 |
| Date | 1 | 0.5388 | 0.5388 | 73.6001 | <0.001 |
| Date$^2$ | 1 | 0.8561 | 0.8561 | 116.9510 | <0.001 |
| Nutrients | 2 | 0.0206 | 0.0103 | 1.4096 | 0.4150 |
| Date * Nutrients | 2 | 0.0124 | 0.0062 | 0.8462 | 0.4314 |
| Date$^2$ * Nutrients | 2 | 0.0206 | 0.0103 | 1.4069 | 0.4155 |

**Table S9.** PERMANOVA of Bray-Curtis dissimilarity between the complete communities with sampling depth as a fixed effect and grouped by plot nested within block.

= Bray-Curtis ~ Sampling Depth + Date * Nutrients, group = Block / Plot

| **Variable** | **df** | **Sum of Sqs** | **R2** | **F statistic** | **p-value** |
|---|---|---|---|---|---|
| Sampling depth | 1 | 1.2344 | 0.0716 | 16.4781 | < 0.001 |
| Date | 1 | 3.6678 | 0.2128 | 48.9609 | < 0.001 |
| Nutrients | 2 | 0.3421 | 0.0198 | 2.2831 | < 0.001 |
| Date * Nutrients | 2 | 0.2283 | 0.0132 | 1.5235 | 0.083 |
| Residual | 157 | 11.7612 | 0.6824 | | |
| Total | 163 | 17.2337 | 1.00000 | | |

**Table S10.** Linear mixed effect model of distance to centroid (dispersion) of Bray-Curtis dissimilarity of the complete communities with sampling depth as a fixed effect and plot nested within block as random intercepts and an ANOVA posthoc test (marginal $R^2 = 0.34$, conditional $R^2 = 0.37$).

= log(Distance to centroid of complete) ~ Sampling Depth + Date * Nutrients + (1 | Block / Plot)

– ANOVA

| Variable | df | Sum of Sqs | Mean Sq | F statistic | p-value |
|---|---|---|---|---|---|
| Sampling Depth | 1 | 0.0358 | 0.0358 | 16.457 | <0.001 |
| Date | 1 | 0.1498 | 0.1498 | 68.688 | <0.001 |
| Nutrients | 2 | 0.1427 | 0.0714 | 65.462 | <0.001 |
| Date * Nutrients | 2 | 0.1412 | 0.0706 | 32.384 | <0.001 |

**Table S11.** PERMANOVA of Bray-Curtis dissimilarity between the core communities with sampling depth as a fixed effect and grouped by plot nested within block.

= Core Bray-Curtis ~ Sampling depth + Date * Nutrients, group = Block / Plot

| Variable | df | Sum of Sqs | R2 | F statistic | P value |
|---|---|---|---|---|---|
| Sampling depth | 1 | 0.9829 | 0.1422 | 45.7364 | < 0.001 |
| Date | 1 | 2.4329 | 0.3520 | 113.2121 | < 0.001 |
| Nutrients | 2 | 0.0618 | 0.0089 | 1.4375 | 0.020 |
| Date * Nutrients | 2 | 0.0602 | 0.0087 | 1.4018 | 0.268 |
| Residual | 157 | 3.3739 | 0.4881 | | |
| Total | 163 | 6.9117 | 1.0000 | | |

**Table S12.** Linear mixed effect model of distance to centroid (dispersion) of Bray-Curtis dissimilarity of the core communities with sampling depth as a fixed effect and plot nested within block as random intercepts and an ANOVA posthoc test (marginal $R^2 = 0.03$, conditional $R^2 = 0.47$).

= log(Distance to centroid for core) ~ Sampling Depth + Date * Nutrients + (1 | Block / Plot) – ANOVA posthoc

| Variable | df | Sum of Sqs | Mean Sq | F statistic | P-value |
|---|---|---|---|---|---|
| Sampling Depth | 1 | 0.1090 | 0.1090 | 6.0187 | 0.0152 |
| Date | 1 | 0.1870 | 0.1870 | 10.3269 | 0.0016 |
| Nutrients | 2 | 0.1636 | 0.0818 | 9.0383 | 0.0031 |
| Date * Nutrients | 2 | 0.1932 | 0.0966 | 5.3346 | 0.0057 |

**Table S13.** Feature scores for survey date and nutrients. Positive features scores are for early survey data and unfertilized plots. Multiple ASVs associated with a taxon are not listed. Feature scores are color coded with blue being the most positive and orange the most negative.
