## Supplementary figures and images for "Seasonal assembly of the phyllosphere fungal microbiome of a perennial grass is robust to nutrient addition"

### Figure S5.pdf

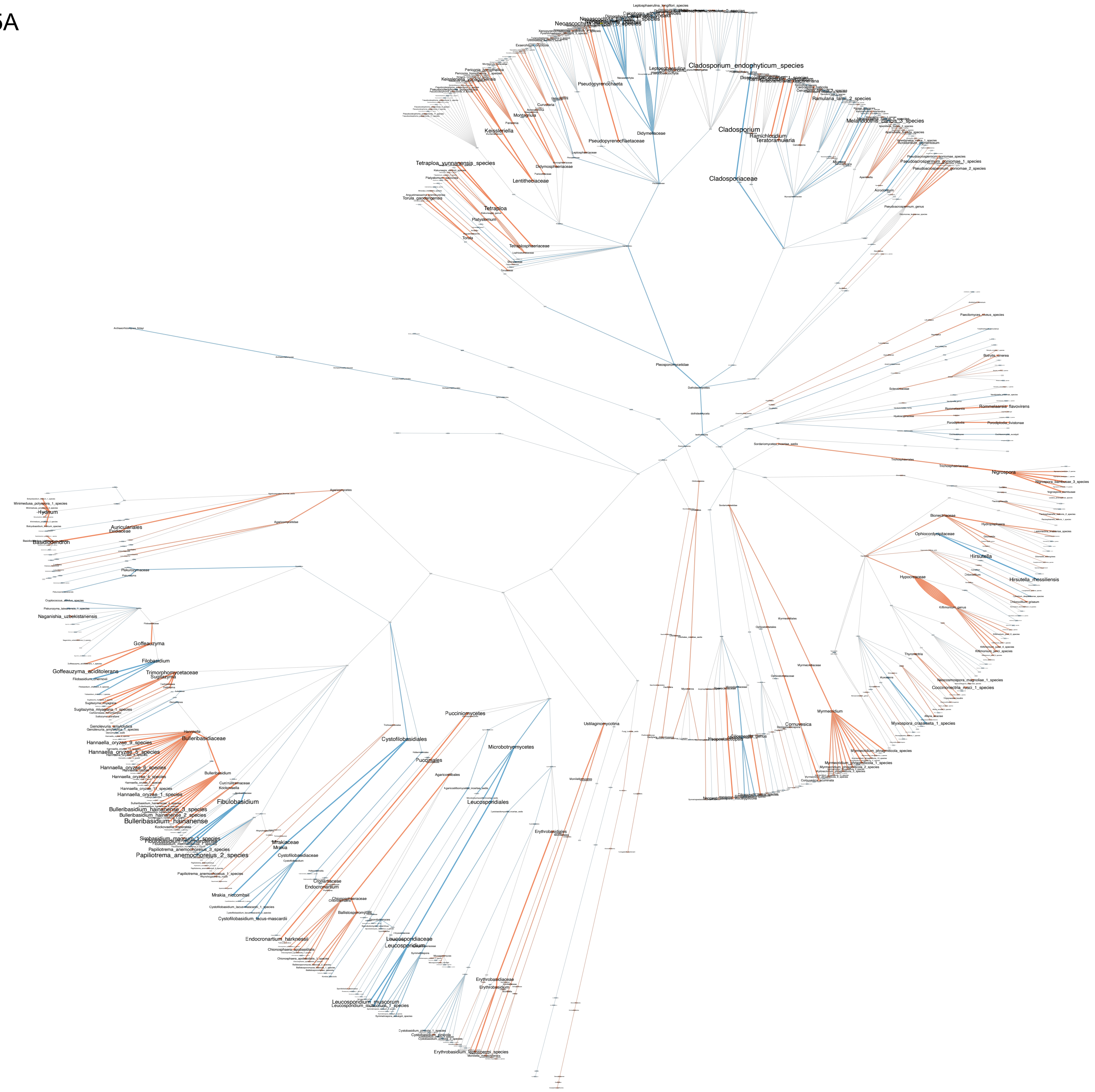

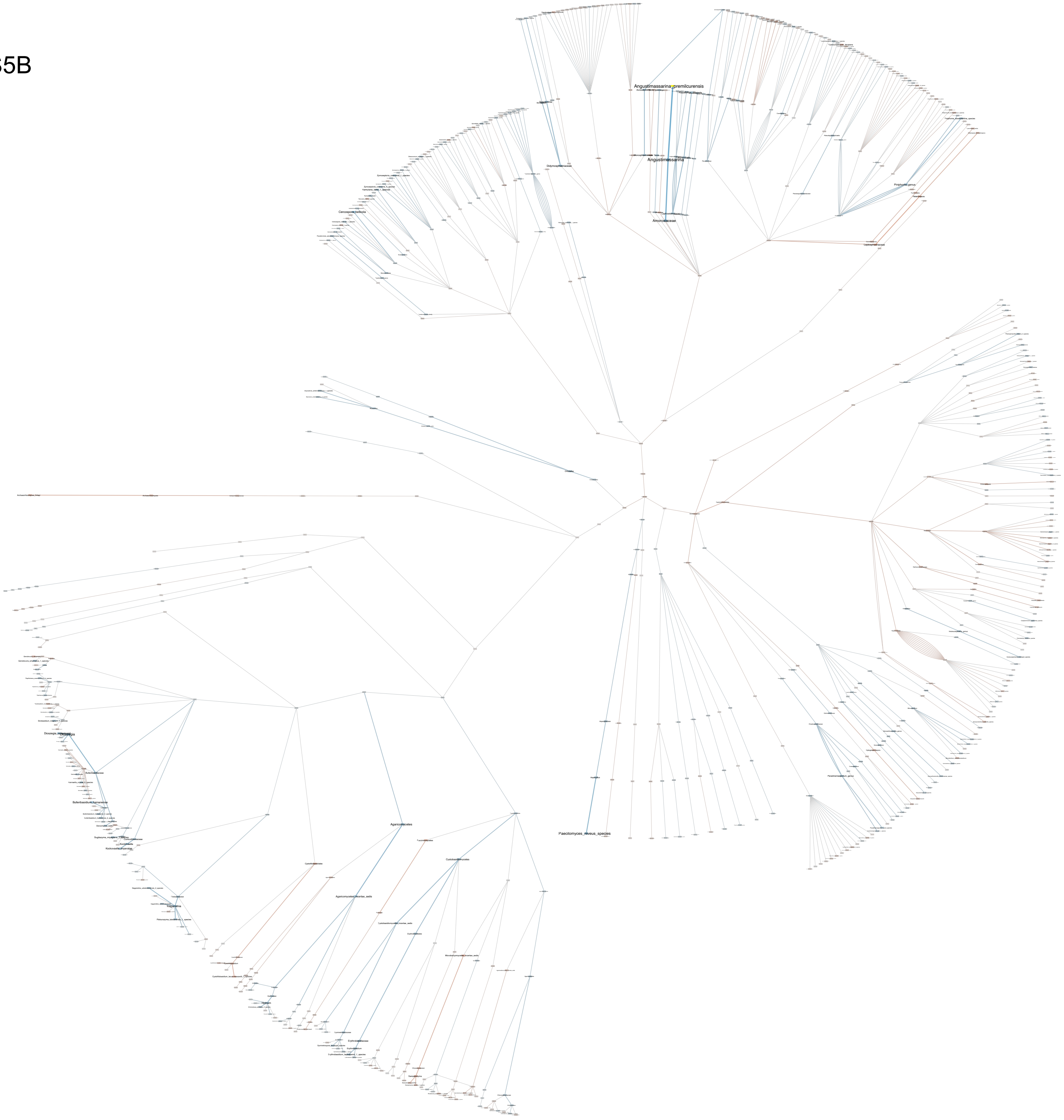
